# RERE and the Mediator complex cooperate with EWSR1::FLI1 in the reprogramming of Translation and Alternative Splicing, the latter being a therapeutically targetable vulnerability in Ewing sarcoma

**DOI:** 10.64898/2026.08.13.744586

**Authors:** Irene Cuervas, Sophie Bonnal, Evelyn Andrades, Silvia Mateo-Lozano, María Sánchez-Jiménez, Pau Berenguer-Molins, Ariadna Acedo-Terrrades, Marta Bódalo-Torruella, Júlia Perera-Bel, Ramón Gimeno, Mònica Roldán, Estela Prada, Juan Valcárcel, Jaume Mora, Inmaculada Hernández-Muñoz

**Author notes:** Corresponding authors: Inmaculada Hernández-Muñoz, PhD Hospital del Mar Medical Research Institute Barcelona Biomedical Research Park (PRBB) C/ Doctor Aiguader, 8808003 Barcelona, Spain, Jaume Mora, MD PhD Pediatric Cancer Center Barcelona (PCCB) Hospital Sant Joan de Déu Passeig Sant Joan de Déu, 2 08950 Barcelona, Spain. Co-senior authors (Jaume Mora, Inmaculada Hernández-Muñoz).

## Abstract

Ewing Sarcoma (ES) is an aggressive neoplasm arising in bones and soft tissues driven by the oncogenic fusion EWSR1::FLI1. Through epigenetic deregulation, EWSR1::FLI1 generates de novo super-enhancers that control the expression of key genes for tumor cell maintenance. By an integrative in silico analysis, we identified the subunit of the Mediator complex MED13L and RERE, a member of the atrophin family of arginine-glutamic acid dipeptide repeat-containing proteins, as genes regulated by EWSR1::FLI1-bound super-enhancers. We confirmed that EWSR1::FLI1 regulates MED13L and RERE expression in ES cell lines and showed that these proteins are highly expressed in Ewing primary tumors. Besides the well-established role of the Mediator complex in transcriptional regulation given its association with the RNA polymerase II, in ES cells the DNA binding sites of MED13L overlap with those of RERE and EWSR1::FLI1 in genes that control protein translation and alternative splicing (AS). Accordingly, the expression of various spliceosome components is co-regulated by MED13L, RERE and the oncogene, leading to AS in ES cells. We identified RBM39, a splicing factor downregulated after MED13L and RERE depletion, as a direct transcriptional target of EWSR1::FLI1. Consistently, *in vitro* viability experiments using indisulam, which induces selective DCAF15-dependent proteosome degradation of RBM39, demonstrate ES cells highly and specifically sensitive to RBM39 inhibition. *In vivo* experiments with mice xenografted with ES cells show complete tumor regression with indisulam, highlighting the potential of this approach as a novel and promising therapeutic strategy for Ewing sarcoma.

**STATEMENT OF SIGNIFICANCE:** Ewing sarcoma (ES) is characterized by FET::ETS oncoproteins that act as pioneer transcription factors. Here, we identified two genes controlled by EWSR1::FLI1-bound super-enhancers, MED13L and RERE, and characterized the mechanism by which these proteins cooperate with the oncogene to regulate RNA metabolism and ribosomal processes in ES cells. These findings have led to the identification of the splicing factor RBM39 as a vulnerability in ES, as supported by the extraordinary sensitivity of these tumors to monotherapy with RBM39 degrader indisulam.

## INTRODUCTION

Ewing sarcoma (ES) is a highly aggressive malignancy arising in bones and soft tissues that predominantly affect children and young adults. Worldwide, it represents the second most common malignant bone cancer in this age group and is the most frequent in Spain^1^. At the molecular level, ES is characterized by a very low mutation burden (0.15 mutations/megabase) and is defined by a pathognomonic chromosomal translocation involving a member of the *FET* gene family and an *ETS* family transcription factor, being *EWSR1::FLI1* the most common translocation (approximately 85% of cases)^2–4^. Importantly, EWSR1::FLI1 can confer transformative capacity on its own when expressed in human embryonic mesenchymal stem cells (heMSCs)^5^. Besides the oncogene, few genetic alterations are observed in ES, such as recurrent loss-of-function mutations in STAG2, a core component of the cohesin-chromatin complex, in 15-20% of patients, which are associated with poor clinical outcomes, including dissemination and accelerated disease progression in advanced stages^3, 4, 6, 7^.

EWSR1::FLI1 acts as a pioneer transcription factor promoting the establishment of lineage-specific *de novo* enhancers and super-enhancers that will interact with their corresponding promoters, leading to the expression of its specific transcriptional program in cells with the permissive cellular context^3, 6, 8–12^. Indeed, EWSR1::FLI1 multimers bind to GGAA microsatellites repeats and recruit histone modifying enzymes and chromatin remodelers, including the SWI/SNF complex, p300, and the MLL complex, thereby converting chromatin regions that would otherwise be closed into regions of euchromatin accessible for transcription^3, 10, 13–17^. In addition, a fraction of EWSR1::FLI1 is indirectly associated with BRD4 at promoter-proximal regions in a complex that includes RNA polymerase II and MED1^18^, the latter being a subunit of the Mediator complex. Mediator is a large multiprotein assembly that communicates regulatory signals from DNA-bound transcription factors directly to RNA pol II, masterfully coordinating development and cell lineage specificity^9, 19, 20^.

In addition to its ability to function as an aberrant transcription factor, EWSR1::FLI1 retains the capacity of the FET proteins to associate with RNA-binding proteins implicated in RNA metabolism^2, 21^. Indeed, 43% of the interactions of the oncoprotein correspond to proteins involved in RNA splicing and processing and 29% with members of the spliceosome complex^22^. For instance, EWSR1::FLI1 binds to the splicing factor SF1 and the small nuclear RNA (snRNA) U1^23, 24^, and to RNA helicase A (RHA), a protein implicated in RNA metabolism whose activity is reduced by the oncogene^25, 26^. Interestingly, the oncogene induces aberrant alternative splicing (AS) of different genes involved in tumorigenesis, such as the SWI/SNF complex protein ARID1A^3, 17, 22, 27^.

In this study, we have characterized the expression and function of two genes, *MED13L* and *RERE*, regulated by EWSR1::FLI1-bound super-enhancers. *MED13L* encodes a subunit of the kinase module of Mediator complex, which can function either as transcriptional activator or repressor depending on the regulatory context^9, 19, 20, 28, 29^. *RERE* (Arginine-Glutamic Acid Dipeptide Repeats) is a nuclear coregulator with poorly characterized functions^30, 31^. Here we report MED13L and RERE being highly expressed in ES, predominantly bound to chromatin regions occupied by other subunits of the Mediator complex and STAG2, and acting as regulators of the expression of numerous spliceosome components, a subset of which are direct oncogene targets, like RBM39. ES cells show high sensitivity to indisulam, an aryl sulfonamide targeting RBM39 by binding as substrate to the E3 ubiquitin ligase DCAF15, demonstrating the critical vulnerability of ES cells to specific AS regulators. Having dissected the role of the Mediator complex and RERE in the splicing control process of ES, we here provide broad *in vivo* evidence for a preclinically highly active new target treatment with RBM39 degrader indisulam, a drug currently in phase II clinical trials^32^.

## RESULTS

### EWSR1::FLI1 binds to *MED13L* and *RERE* super-enhancers in Ewing sarcoma cells

To identify genes regulated by super-enhancers that might play a role in ES tumorigenesis, we crossed three databases: super-enhancer-regulated genes bound by EWSR1::FLI1 in the A673 cell line^11^; the transcriptional profiles of ES tumors^5^ and the transcriptome of murine ES generated from embryonic chondrogenic progenitors expressing EWSR1::FLI1^33^. Forty-five common genes were identified, and among them, *MED13L*, one of the Mediator complex subunits, and the transcriptional regulator *RERE* were selected (**Figure 1A, S1A, S1B**). MED13L is a transcriptional activator as well as repressor in important neural and cardiac developmental pathways and reported to be involved in the recruitment of p300 to enhancers and promoters in non-small-cell-lung cancer^9, 19, 20, 28, 29, 34–36^. RERE is a nuclear coregulator implicated in embryonic development^30, 31^. Deficiency of RERE results in a phenotype that overlaps with that observed in individuals with CHARGE syndrome, which is associated with a deficiency in CHD7 (chromodomain helicase DNA binding protein 7), a chromatin remodeler involved in the formation of DNA loops at super-enhancers^37–40^. Expression data extracted from DepMap (https://depmap.org/portal) show RERE mRNA levels as significantly higher in ES cell lines and MED13L mRNA levels similar in all cell lines (**Figure 1B)**. Likewise, mRNA levels of RERE, but not those of MED13L, are higher in ES tumors compared to other pediatric tumor types (**Figure S1C**). However, analysis of protein expression by immunohistochemistry in primary ES, rhabdomyosarcoma and neuroblastoma tissues showed both MED13L and RERE specific and significantly higher in ES (**Figure 1C**). Other Mediator subunits such as MED1 and MED12 were equally expressed among the three developmental tumors analyzed (**Figure 1C**).

**Figure 1.**
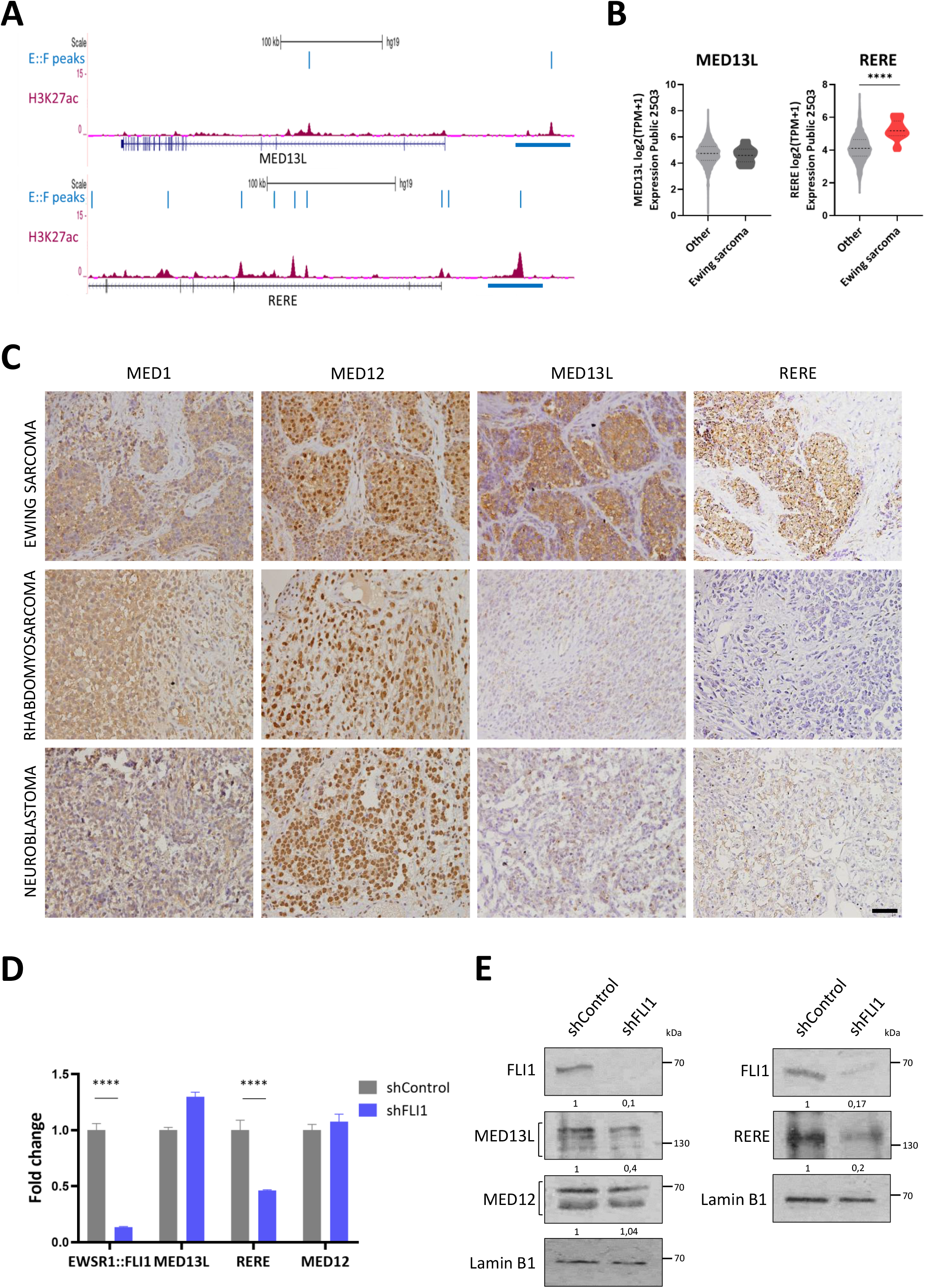
MED13L and RERE are differentially expressed in Ewing sarcoma tumors. **A)** Tracks from the University of California Santa Cruz (UCSC) Genome Browser (https://genome.ucsc.edu/) showing H3K27ac enrichment and peaks corresponding to EWSR1::FLI1 binding in MED13L (top, chromosome 12) and RERE (bottom, chromosome 1) in A673 cells11. Blue bars, super-enhancer regions with EWSR1::FLI1 binding. E::F, EWSR1::FLI1. **B)** mRNA expression of MED13L and RERE in cell lines from adult and pediatric tumors (n=1576) compared to Ewing sarcoma cell lines (n=24), extracted from the Expression Public 25Q3 database from the Cancer Dependency Map (https://depmap.org). Unpaired t test by the Mann Whitney test was performed to evaluate the differences. ****P <0.0001. **C)** Immunohistochemistry (IHC) for MED1, MED12, MED13L and RERE expression in representative samples of Ewing sarcoma, rhabdomyosarcoma and neuroblastoma tumors obtained from Hospital Sant Joan de Déu Barcelona Biobank. Black bar, scale bar: 50 μm. **D)** mRNA expression levels for EWSR1::FLI1, MED12, MED13L and RERE in A673 cells treated with 1 μg/mL doxycycline to induce shFLI1 expression and determined by RT-qPCR. Error bars indicate the standard deviation of three replicates. Values were normalized to Actin B. Two-way ANOVA was used to analyze differences and Sidak’s multiple comparisons test was performed to compare expression versus control. ****P value < 0.0001. **E)** Expression levels of MED12 and MED13L (whole-cell lysate) and RERE (nuclear extract) in A673 cells after inducing EWSR1::FLI1 knockdown with 1 μg/mL of doxycycline, determined by Western blot (WB). Lamin B1, loading control. Square brackets indicate the protein isoforms detected. Quantification of band intensities relative to Lamin B1 was performed with Image J software.

To determine whether EWSR1::FLI1 is required for MED13L and RERE sustained expression, we took advantage of the ES cell line A673 carrying a doxycycline-inducible short hairpin RNA (shRNA) targeting FLI1^41^. Downregulation of the oncogene expression resulted in a decrease of RERE mRNA levels while MED13L and MED12 mRNAs remained unaffected (**Figure 1D**). In contrast, EWSR1::FLI1 silencing reduced MED13L and RERE protein levels by more than 50%, whereas MED12 was unaffected (**Figure 1E**). Taken together, these results confirm that expression of RERE depends on EWSR1::FLI1, which is consistent with its identification as a gene transcriptionally regulated by the binding of EWSR1::FLI1 to its super-enhancer, while the reduction at the protein, but not mRNA, level of MED13L upon oncogene silencing suggests a posttranscriptional mechanism which could involve the expression of antisense MED13L non-coding transcripts.

### MED13L and RERE preferentially bind to common intergenic regions in Ewing sarcoma cells

The Mediator complex binds to many activator-bound enhancer regions and promoters, facilitating the activation of gene transcription^42^. As for RERE, the similarity of the phenotype between the deficiency of CHD7^37–40^ and RERE suggested that they might share some chromatin-binding abilities. We explored the chromatin regions bound by MED13L and RERE in ES cells, and those of MED1 and MED12 subunits, whose expression is not regulated by the oncogene, by chromatin immunoprecipitation sequencing (ChIP-seq). In addition, we determined chromatin regions enriched in acetylated histone lysine 27 (H3K27ac), which characterizes enhancers, super-enhancers and promoters. The chromatin sites bound by EWSR1::FLI1 were obtained from Bilke *et al*.^43^ and were used to identify the regions shared by Mediator, RERE and the oncogene in A673 cells.

Genomic distribution of ChIP-seq peaks showed that H3K27ac and EWSR1::FLI1 signals were enriched at promoter regions, as expected (**Figure 2A**). In contrast, peaks of the Mediator subunits and RERE were mostly found in distal intergenic regions. With respect to their binding to gene body regions, MED1, MED13L and RERE displayed a preferential enrichment for introns, while EWSR1::FLI1, H3K27ac and MED12 were equally enriched in intronic and exonic regions. These data suggested that genome-wide distribution of MED13L and RERE differs from EWSR1::FLI1, which is further supported by differences in the DNA-binding motifs found in MED13L and RERE-enriched regions: the oncogene binding sites are enriched in well-described GGAA repetitive motifs^10, 13–16^, whereas MED13L is preferentially bound to GGAAT motifs in distal intergenic regions (**Figure 2B**), which correspond to the core binding site of TEA domain (TEAD) family of transcription factors^44^. Binding of MED13L and RERE to intronic, exonic and promoter regions, and to distal intergenic regions for RERE, was preferentially found in CA microsatellite regions (**Figure 2B**), which cooperate in the regulation of gene expression by establishing a left-handed DNA (Z-DNA) conformation^45^.

**Figure 2.**
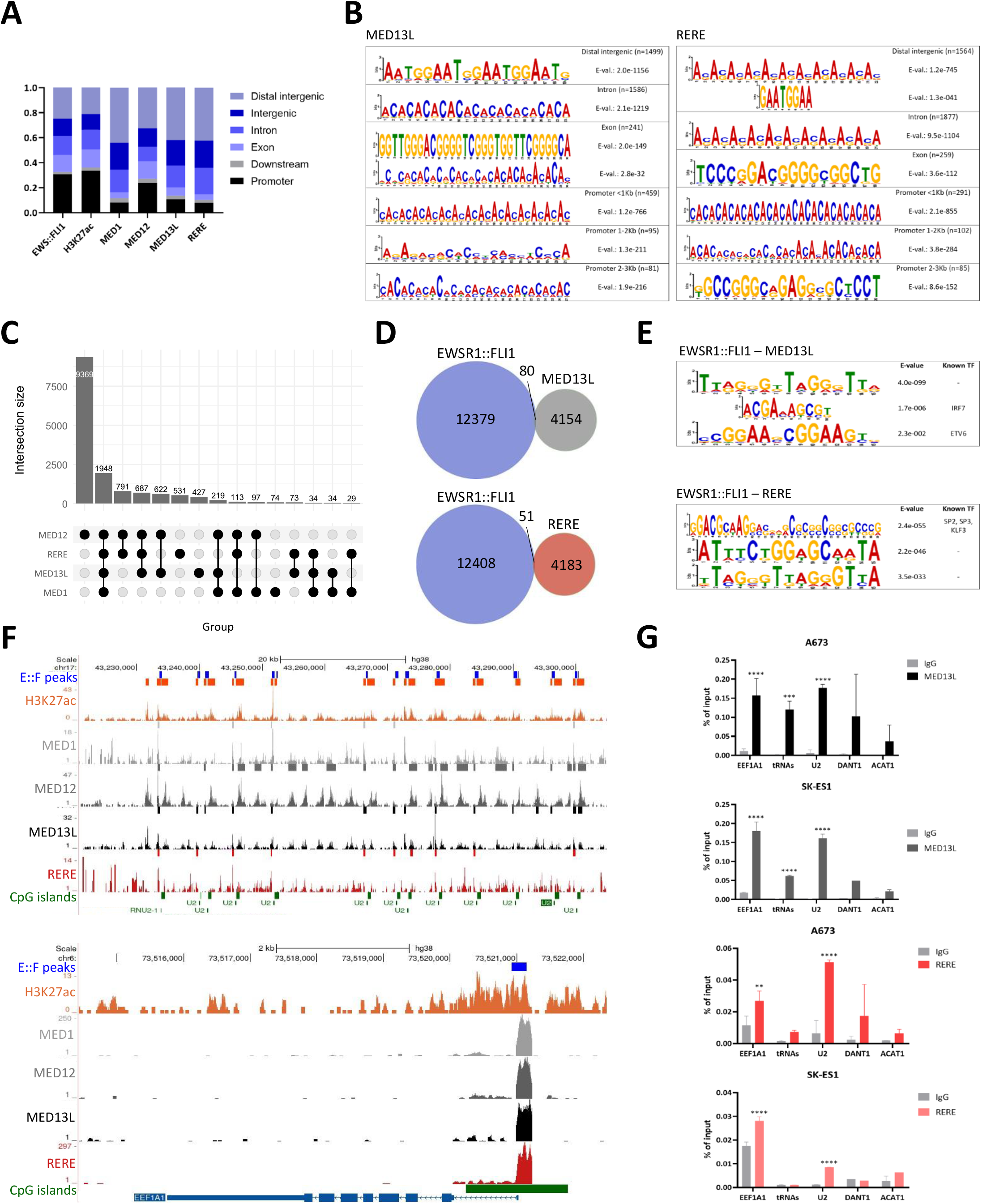
Genome-wide binding of MED13L and RERE largely differs from EWSR1::FLI1 binding but overlaps in a subset of chromatin regions. **A)** Bar plots depicting the genomic distribution of EWSR1::FLI143, H3K27ac, MED1, MED12, MED13L and RERE in A673 cells. **B)** Top MEME DNA binding motifs for MED13L and RERE peaks and their corresponding E value in distal intergenic, intronic, exonic and promoter regions. **C)** Upset plot showing the intersection of ChIP-seq peaks for MED1, MED12, MED13L and RERE in A673 cells. **D)** Venn diagrams showing the overlap between ChIP-seq peaks of MED13L or RERE and EWSR1::FLI143 in A673 cells. **E)** MEME DNA binding motifs for overlapping peaks in the same genomic regions of EWSR1::FLI143 with MED13L or RERE and their corresponding E value. TF, transcription factor. **F)** UCSC Genome Browser tracks for the binding of EWSR1::FLI143, MED1, MED12, MED13L and RERE, as well as H3K27ac enrichment in the snRNA U2 (top) and the EEF1A1 genic regions (bottom) (in chromosomes 17 and 6, respectively) in the A673 cell line. E::F, EWSR1::FLI1. **G)** Validation by ChIP-qPCR of MED13L and RERE binding to targets overlapping with EWSR1::FLI1 in the Ewing sarcoma cell lines A673 and SK-ES1. Values represent the enrichment ratio of immunoprecipitated samples relative to input. IgG, negative control for non-specific binding; ACAT1, negative control region. Two-way ANOVA was used to analyze differences and Sidak’s multiple comparisons test was performed to compare enrichment versus IgG. **P value=0.0031, ***P value=0.0007, ****P value <0.0001.

To date, no relationship between Mediator and RERE has been reported. Since our ChIP-seq analysis revealed a striking similarity in the genomic distribution and the underlying consensus DNA sequences of RERE and some Mediator subunits, we intersected the binding peaks of each protein to elucidate whether they bind to the same chromatin regions. Eighty per cent of the RERE peaks (corresponding to 3876 peaks) overlapped with peaks bound by at least one of the three Mediator subunits analyzed (MED1, MED12 and MED13L), and 44% of the RERE peaks (1948 peaks) overlapped with all three units simultaneously (**Figure 2C**). These data indicated that RERE mainly binds to the same chromatin regions as the Mediator complex, which are specific for ES cells, suggesting that they may functionally cooperate.

### MED13L and RERE bind to a subset of EWSR1::FLI1-target genes involved in RNA metabolism and ribosomal processes

Our ChIP-seq analyses showed that the distribution of chromatin regions enriched in MED13L and RERE largely differed from that observed in EWSR1::FLI1-binding regions. To determine whether there were chromatin sites where the oncogene coincided with MED13L and RERE recruitment, we intersected MED13L and RERE peaks with those of EWSR1::FLI1^43^. Consistent with the divergent genomic distribution of the peaks, this analysis identified only 80 and 51 common peaks with MED13L and RERE, respectively (**Figure 2D**). Gene ontology (GO) of this subset of genes corresponded to ribosome structure and function (**Figure S2A**). Analysis of the consensus DNA-binding motif for these regions identified the IRF7 motif and the GGAA single ETV6 motif for MED13L, and SP2, SP3 and KLF3 binding motifs for RERE (**Figure 2E**). These motifs are distinct from GGAA microsatellite repeats that are specifically bound by EWSR1::FLI1 within *cis*-regulatory regions^3, 10, 13–17^.

The discrete MED13L, RERE and oncogene overlapping regions were mainly characterized by their enrichment in CpG islands. Most of these peaks were distributed in tandem and corresponded to non-protein coding regions. Examples of the tandem overlapping regions include, among others, the snRNA *U2*, an essential component of the splicing machinery that is organized as a nearly perfect tandem array containing 5 to 22 copies of a 5.8 kb repeated unit^46^ (**Figure 2F**, top); the ribosomal subunits *RNA5S* and *RNA5.8S*; the *tRNA* loci, directly involved in ribosome function (**Figure S2B**); and *DANT1*, a long non-coding RNA with a role in X chromosome inactivation in female cells. Notably, the genomic region corresponding to *EEF1A1*, which encodes a protein responsible for the enzymatic delivery of aminoacyl tRNAs to the ribosome, also represented a co-bound overlapping region regulated by CpG islands (**Figure 2F**, bottom). **Figure 2G** shows specific MED13L and RERE binding to some of these genomic regions in ES, but not in rhabdomyosarcoma cells (**Figure S2C**), as determined by ChIP-qPCR.

### MED13L, RERE and STAG2 co-bind to many EWSR1::FLI1-target genes

Genome browser (https://genome.ucsc.edu/) visualization of the chromatin regions enriched in Mediator and RERE and the EWSR1::FLI1 binding peaks showed that, although most of the peaks of these two proteins did not overlap with those of the oncogene, they were often located in the same genes. Crossing of genes associated with MED13L and RERE peaks and those associated with EWSR1::FLI1 revealed that 34% of MED13L-bound genes (1108 genes) and 34% of RERE-bound genes (1160) were coincident with genes associated with EWSR1::FLI1 peaks (**Figure 3A**). An example of a gene co-occupied by Mediator, RERE and EWSR1::FLI1 is *LINGO1*, as shown in **Figure 3B**. In this genomic region, EWSR1::FLI1 binds to a distal intergenic region, whereas Mediator and RERE are bound to the gene promoter.

**Figure 3.**
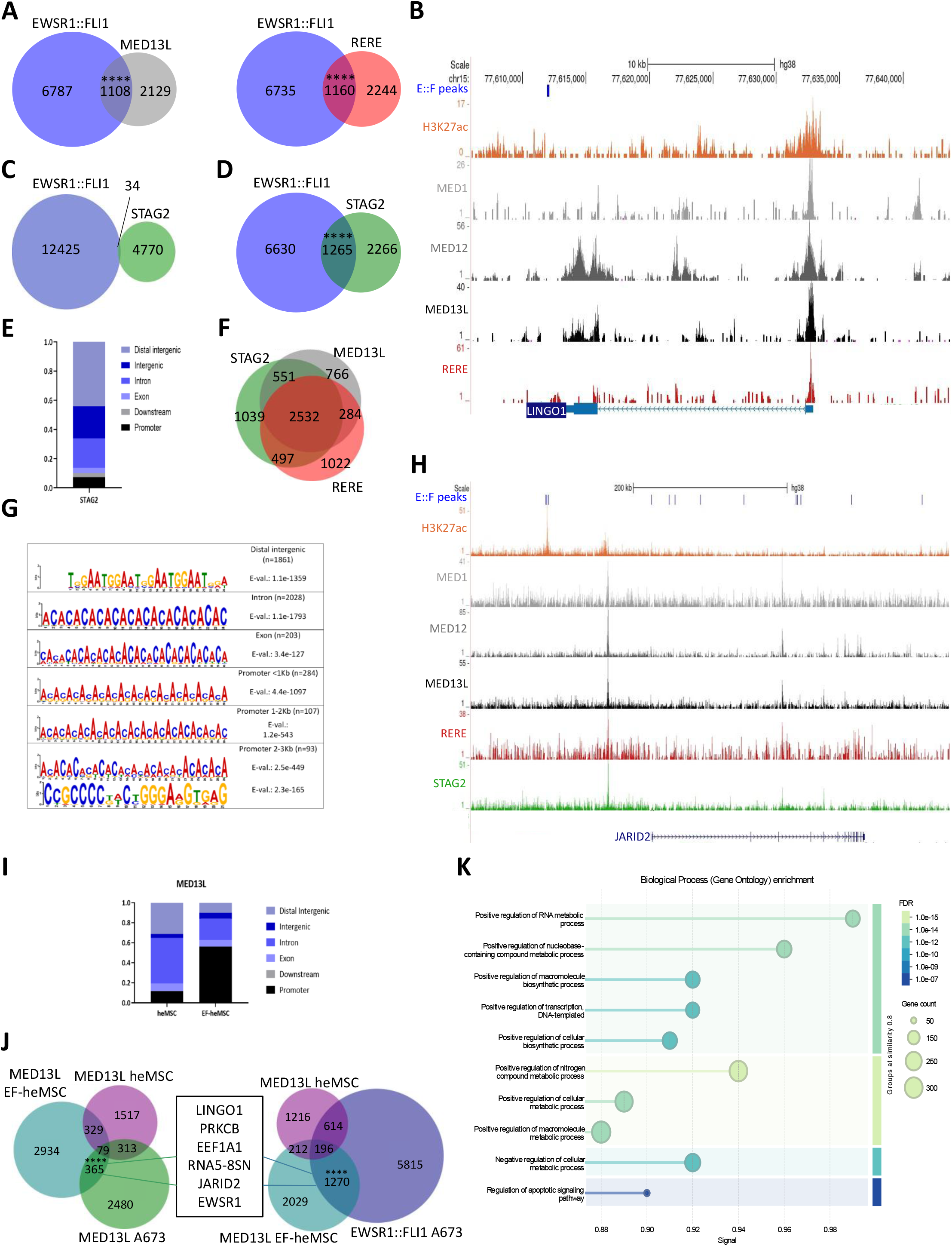
MED13L, RERE and STAG2 binding to EWSR1::FLI1-target genes. **A)** Intersection between genes associated with MED13L and RERE binding peaks with genes associated with EWSR1::FLI1 binding peaks43 in A673 cells. Fisher’s exact test was used to statistically evaluate the association of each gene set. Odds ratio for MED13L and RERE with EWSR1::FLI1 was 1.8. ****P <0.0001. **B)** ChIP-seq signal tracks for FLI143, H3K27ac, MED1, MED12, MED13L and RERE in the LINGO1 genic region, generated with the UCSC Genome Browser. **C)** Venn diagram depicting STAG2 and EWSR1::FLI1 overlapping peaks in A673 cells. **D)** Intersection of genes associated with STAG2 and EWSR1::FLI1 binding peaks43 in A673 cells. Fisher’s exact test was used to statistically evaluate the association of each gene set. Odds ratio for STAG2 with the oncogene was 2. ****P <0.0001. **E)** Bar plots depicting STAG2 genomic distribution in A673 cells. **F)** Venn diagram showing the intersection of MED13L, RERE and STAG2 peaks in A673. **G)** MEME DNA binding motif for STAG2 peaks and their corresponding E value in distal intergenic, intronic, exonic and promoter regions. **H)** UCSC Genome Browser tracks showing the enrichment of FLI143, H3K27ac, MED1, MED13L, RERE and STAG2 in the JARID2 genomic region (chromosome 6). E::F, EWSR1::FLI1. **I)** Genomic distribution of MED13L in heMSC cells with absence (heMSC) or presence of the oncogene (EF-heMSC), depicted in bar plots. **J)** Left panel, Venn diagram showing the intersection of genes associated with MED13L peaks in A673, heMSC and EWSR1::FLI1-heMSC cells. Fisher’s exact test was used to evaluate the association of EF-heMSC and A673 conditions. Odds ratio corresponds to 1.4. ****P <0.0001. Right panel, intersection of genes associated with EWSR1::FLI1 peaks in A673 cells43 and those associated with MED13L peaks in EWSR1::FLI1-heMSC cells. Odds ratio after Fisher’s exact test for overlapping genes in A673 and EF-heMSC cells, 2.7. ****P <0.0001. **K)** Gene ontology enrichment of the genes identified in I, performed with STRING88 (https://string-db.org/). FDR, False Discovery Rate.

The cohesin member STAG2 promotes interactions between enhancers and promoters, and the oncogenic program driven by EWSR1::FLI1 is highly perturbed in STAG2 knockout cells, due in part to altered enhancer-promoter contacts^42, 47, 48^. Therefore, given that one of the canonical functions of Mediator is the formation of DNA loops, we performed ChIP-seq experiments to determine STAG2 binding to chromatin in the A673 cell line and to compare its genome-wide localization to that of the oncogene and Mediator. As described for MED13L and RERE, the overlapping peaks of STAG2 and EWSR1::FLI1 binding were also quite limited, with only 34 common peaks (**Figure 3C**), while the intersection of STAG2 and EWSR1::FLI1-bound genes was similar to that of the oncogene and MED13L and RERE (35%; 1265 genes; **Figure 3D**). As observed for MED13L and RERE genome-wide binding, most STAG2 genomic enrichments corresponded to distal intergenic regions and introns, rather than exons, promoters or super-enhancers (**Figure 3E**). Importantly, 78% of the STAG2-bound regions (3780 peaks) were also enriched in MED13L or RERE (**Figure 3F**), and these three proteins share the DNA binding motifs GGAAT and CACA microsatellites identified for MED13L and RERE (**Figure 3G**). **Figure 3H** shows an example of a gene bound by STAG2 and the oncogene, *JARID2*, along with Mediator and RERE.

Since MED13L and RERE protein levels rely on EWSR1::FLI1 expression, it is not possible to study their dependence on the oncogene for chromatin recruitment in cells in which the oncogene has been depleted. Therefore, we used human embryonic mesenchymal stem cell (heMSCs) in which we have shown that EWSR1::FLI1 expression recapitulates ES transcriptome and tumorigenesis^5^, and we performed ChIP-seq to detect MED13L binding before and after EWSR1::FLI1 expression in these cells. Analysis of the genomic distribution of the peaks revealed that, in the absence of oncogene expression, MED13L occupancy was enriched at intronic regions (**Figure 3I**). In contrast, upon EWSR1::FLI1 expression, MED13L binding sites were predominantly localized to promoter regions, indicating an oncogene-mediated redistribution of MED13L across the genome early in ES.

To identify the changes in MED13L distribution induced by the oncogene in early stages of ES tumorigenesis, we intersected the genes associated with MED13L peaks upon oncogene expression in heMSCs with those genes associated with MED13L peaks in the A673 cell line. This approach showed that the subset of genes bound by MED13L in EWSR1::FLI1-heMSCs (but not in parental heMSCs) and in A673 included genes identified as MED13L targets in ES, such as *LINGO1*, *PRKCB*, *EEF1A1*, *RNA5-8SN*, *JARID2* or *EWSR1* (**Figure 3J**, left). Furthermore, the intersection of genes associated with MED13L peaks in EWSR1::FLI1-heMSCs and genes associated with EWSR1::FLI1 peaks in A673 cells showed a significant overlap including the above mentioned MED13L target genes (**Figure 3J**, right). Gene ontology analysis of these overlapping genes revealed their role in RNA metabolic processes (**Figure 3K**), consistent with the function of genes jointly bound by MED13L, RERE, STAG2 and EWSR1::FLI1 in A673 cells. These results indicate that MED13L redistribution to a subset of its genomic target regions in ES is mediated by EWSR1::FLI1 in early stages of ES tumorigenesis.

### MED13L and STAG2 colocalize with EWSR1::FLI1 in nuclear compartments

Our results showing MED13L, RERE and STAG2 overlapping in a notable proportion of peaks and their co-occupancy in a significant number of oncogene-bound genes suggested that their distribution within the nucleus might be in close proximity. To investigate this possibility, we analyzed their subcellular distribution by immunofluorescence (IF). To overcome technical limitations of MED13L and FLI1 antibodies produced in rabbit, A673 cells were transduced with a FLAG-tagged EWSR1::FLI1 lentiviral supernatant, allowing the detection of the tagged oncogene with a mouse FLAG antibody. Validation of this approach (**Figure S3A**) was followed by simultaneous detection of MED13L and FLAG in transfected cells (GFP positive) (**Figure S3B**). Confocal imaging showed the subcellular localization of MED13L being mainly nuclear, but also exhibited cytoplasmic distribution, while the FLAG signal of the transduced oncogene was solely nuclear, as expected (**Figure 4A)**. To better characterize the IF signals, we recorded the intensity histogram of each fluorochrome in a selected nuclear region. Plot profiling showed that the fluorescent signals of FLAG and MED13L were coincident (**Figure 4A**), suggesting that FLAG-EWSR1::FLI1 and MED13L colocalize in discrete nuclear regions. Immunoprecipitation (IP) assays in A673 extracts with antibodies against FLI1, MED1, MED12 and MED13 confirmed that Mediator subunits interact among themselves as part of the same complex, and MED13L and MED12 immunocomplexes effectively pulled down endogenous EWSR1::FLI1 (**Figure 4B**). Although MED12 and MED13L could not be detected when the oncogene was pulled down with a FLI1 antibody, probably due to steric hindrance effects, cell extracts from A673 cells expressing the FLAG-tagged oncogene were immunoprecipitated with a FLAG antibody and Western blot confirmed the interaction of endogenous MED13L with FLAG-EWSR1::FLI1 (**Figure S3C**).

**Figure 4.**
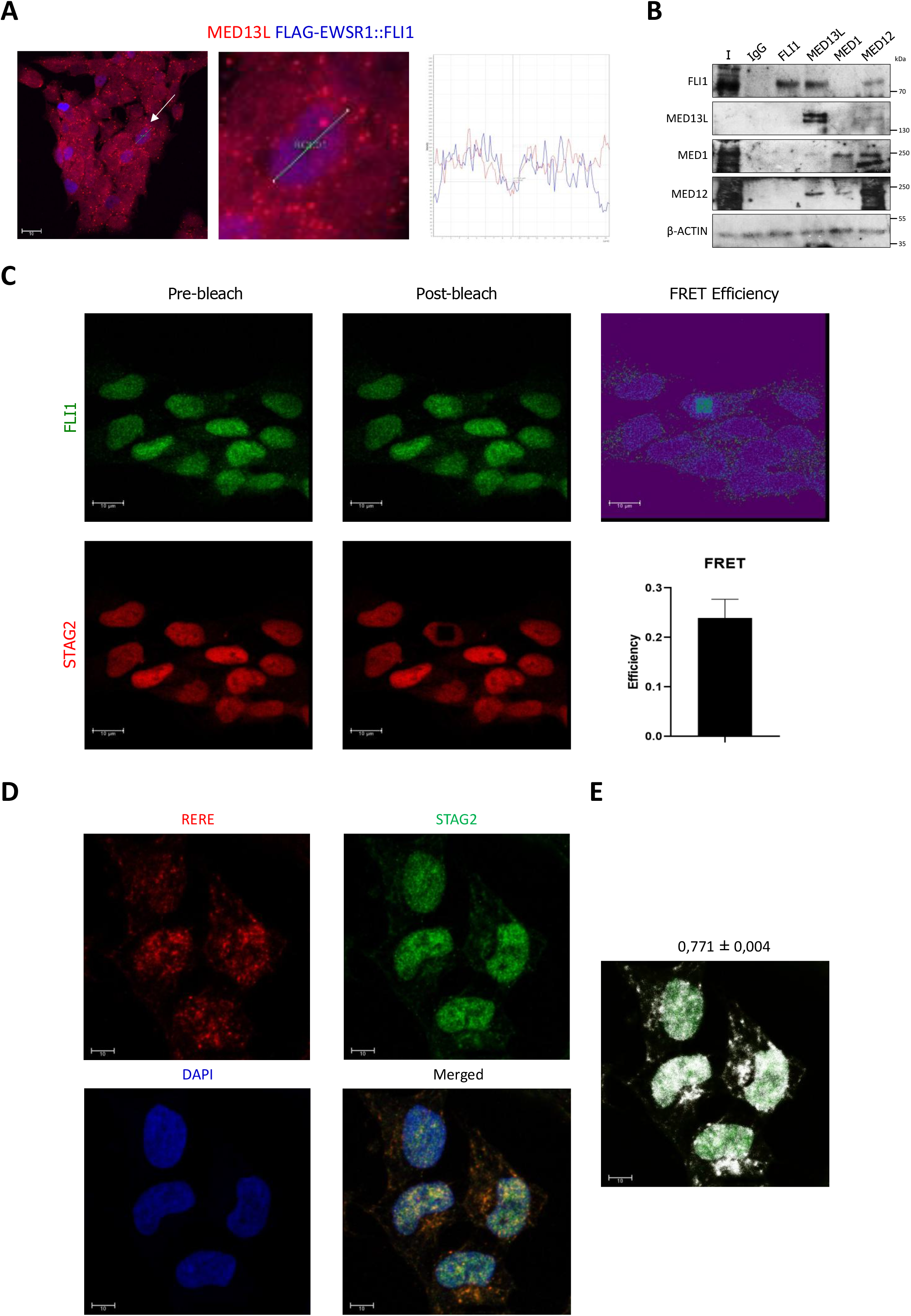
MED13L and STAG2 colocalize with EWSR1::FLI1. **A)** Representative image of endogenous MED13L (red) and ectopic FLAG-EWSR1::FLI1 (blue) in the nuclei of A673 cells, obtained by immunofluorescence (IF). Scale bar: 10 μm. On the right, a magnified image of a single cell and histogram of the intensities of the corresponding MED13L and FLAG signals in this image. **B)** Western blot detection of proteins immunoprecipitated with antibodies for FLI1, MED1, MED12 and MED13L in A673 cell extracts. IgG was used as a negative control. Ten percent of the cell lysates were used as input. **C)** Colocalization analysis of STAG2 and endogenous EWSR1::FLI1 by fluorescence resonance energy transfer (FRET). Left and middle images depict EWSR1::FLI1 and STAG2 immunofluorescence signals before and after photobleaching. Sixteen regions of interest (ROI) were analyzed from different cells. FRET efficiencies were calculated during STAG2 photobleaching. Data represents S.E. **D)** IF staining of STAG2 (green) and RERE (red) in A673 cells. Cell nuclei were stained with DAPI staining (blue). Scale bar: 10μm. **E)** Mask image showing quantification of colocalization by Overlap Coefficient in A673 cells. Scale bar: 10μm. Data represents mean ± S.E. of two fields. Capture of confocal images and signal quantification were performed using the LAS X software (Leica).

We next investigated the subnuclear localization of STAG2 and EWSR1::FLI1 by IF, in this case detecting the endogenous oncoprotein. This analysis revealed that both proteins were in close proximity, less than 100 Å in the nuclei of A673 cells, since the Fluorescence Resonance Energy Transfer (FRET) efficiency of the interaction was 0.24 (**Figure 4C**). In addition, according to RERE and STAG2 genome wide binding to overlapping chromatin regions, RERE and STAG2 colocalized in discrete nuclear domains identified by the presence of yellow aggregates (**Figure 4D**). Confocal quantification of the signals resulted in an Overlap Coefficient correlation of 0.77, indicating a less than 290 nm distance between the two proteins (**Figure 4E**). Altogether, these data indicate that EWSR1::FLI1 colocalizes and interacts with Mediator, while EWSR1::FLI1 and STAG2 are in the same nuclear domains. In addition, STAG2 and RERE are also localized within the same nuclear domains.

### MED13L and RERE regulate the expression of genes involved in RNA metabolism and ribosomal processes

To investigate the functional role of MED13L and RERE, transient depletion by siRNA in A673 cells was performed (**Figure S4A**), and changes in their transcriptomes were determined. RNA sequencing (RNA-seq) and independent analysis of the changes induced by three different siRNAs targeting each of these proteins identified 4,866 differential expressed genes (DEGs; 1,862 and 3,004 down-regulated and up-regulated, respectively; *p* < 0.05) in MED13L-depleted cells and 4,295 DEGs in RERE-depleted cells (1,657 and 2,638 down-and up-regulated respectively; *p* < 0.05). Of note, only a few MED13L-or RERE-bound genes identified by ChIP-seq displayed significant transcriptional changes, accounting for about 6% of the DEGs, suggesting that direct DNA binding is not the main mechanism contributing to the MED13L-or RERE-mediated transcriptional profile in ES cells. Likewise, many of the genes associated with the ES gene signature are not direct targets of EWSR1::FLI1 in A673 cells^11^.

Gene set enrichment analysis (GSEA) of the DEGs showed a reduction in genes coding for proteins involved in RNA metabolism and splicing, as well as in ribosomal processes, in cells with MED13L or RERE depletion (**Figure 5A, 5B**, respectively). On the contrary, MED13L and RERE defective cells presented enhanced expression of genes involved in cell structure and migration processes, and in lipid metabolism (**Figures S4B, S4C**). Since MED13L and RERE bind to overlapping chromatin regions and share the underlying consensus DNA sequences (**Figure 2B, 2C**), we intersected the downregulated transcripts in cells with reduced levels of each of these two proteins. Fifty-three per cent and sixty per cent of the genes whose expression was reduced in cells with MED13L and RERE depletion, respectively, were common to both conditions (corresponding to 996 transcripts) (**Figure S4D**). GO enrichment of these transcripts showed their involvement in RNA splicing and processing (**Figure S4D**). In contrast, GO of the genes whose expression was reduced following MED13L depletion was cytoplasmic translation and ribosome biogenesis, whereas RERE-deficient cells showed a specific reduction in the expression of genes involved in cell cycle (**Figure S4D**). These data indicated that RERE and MED13L functionally cooperate in the transcriptional regulation of RNA splicing and processing.

**Figure 5.**
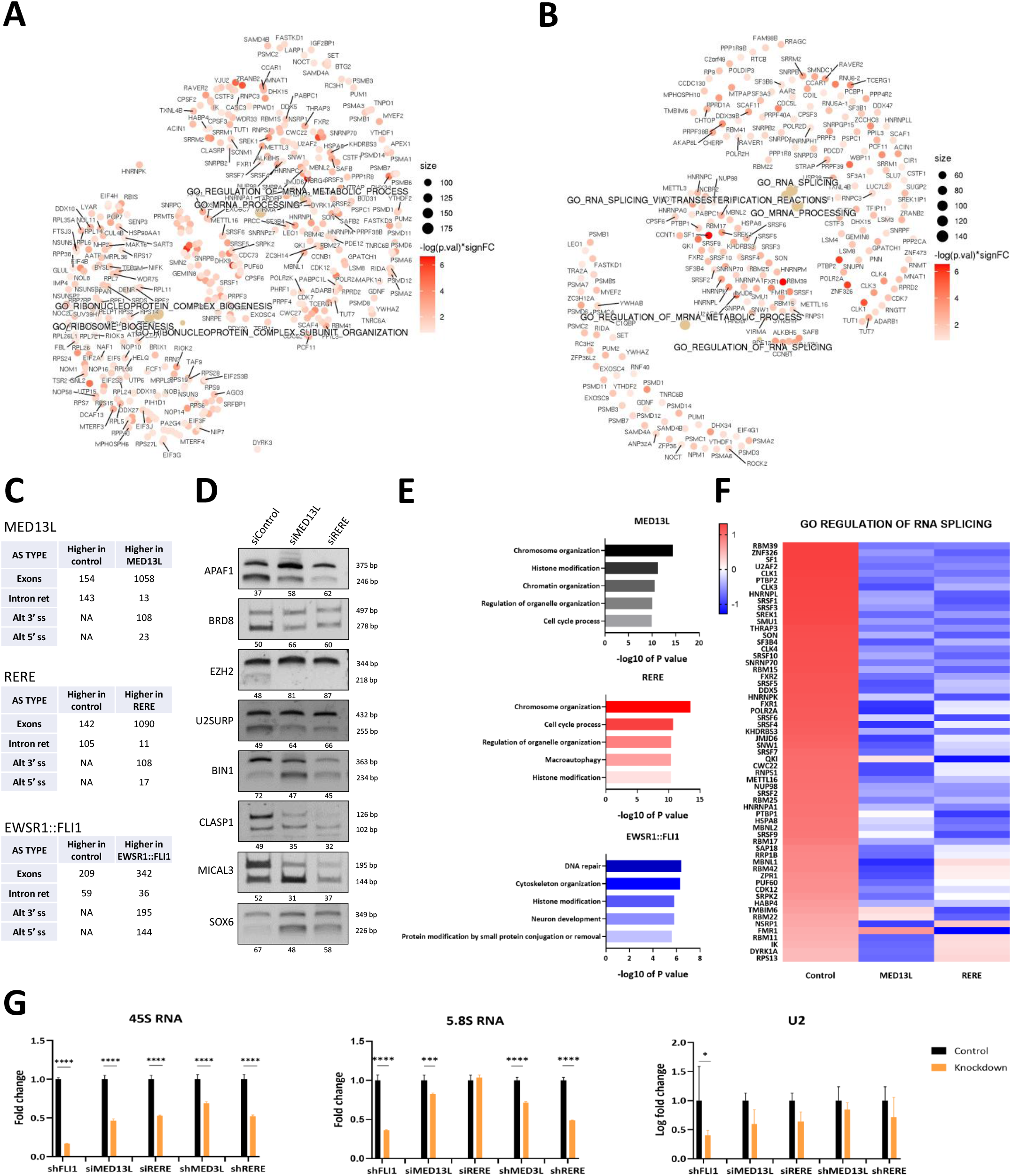
The expression of splicing factors and ribosomal RNAs depends on MED13L and RERE. A, **B)** Gene-Concept Networks of the top five significantly downregulated terms in MED13L **(A)** and RERE **(B)** depleted cells by three different siRNA sequences. The subset of represented genes that contributed most to the enrichment result was ordered by-log(P. value)*significant FC. FC, fold change; NES, normalized enrichment score. **C)** Summary of the VAST-TOOLS output comparing three controls (siControl) vs. three knockdown samples (siMED13L, siRERE) and esiEWSR1::FLI150. The events were classified as exons, intron retention, alternative 3’ splice site (alt. acceptor) and alternative 5’ splice site (alt. donor) events. For each category, the number of events higher in control and higher upon knockdown after applying a dPSI of 15 are shown. **D)** Validation by RT-PCR of alternative splicing changes in selected genes induced by MED3L and RERE depletion by siRNA in A673 cells. Sequences #1 for MED13L and RERE siRNA knockdown were used to perform these experiments. Percent spliced in (PSI), calculated with ImageJ is indicated under each gene **E)** GO biological process categories for 1,191 genes associated to AS events induced by siRNA MED13L knockdown, 1,162 genes associated to AS events induced by RERE knockdown by siRNA, and 846 genes with AS due to EWSR1::FLI1 esiRNA50. Analyses were performed with the ToppGene89 online database (https://toppgene.cchmc.org/). **F)** Heat map depicting significant expression changes of genes involved in mRNA splicing after MED13L and RERE knockdown by siRNA in the A673 cell line. **G)** 45s and 5.8s ribosomal RNA and U2 snRNA expression in A673 cells depleted from FLI1, MED13L and RERE. Values were normalized to Actin B. Two-way ANOVA was used to analyze differences and Sidak’s multiple comparisons test was performed to compare expression versus control. *P value <0.05, ***P value=0.0005, ****P value <0.0001. GO, gene ontology.

To better characterize the role of MED13L and RERE in the transcriptional regulation of AS genes, we carried out an analysis of the RNA-seq data from MED13L-and RERE-depleted cells using the pipeline VAST-TOOLS^49^. In parallel, public RNA-seq database of A673 cells harboring an EWSR1::FLI1-endoribonuclease-prepared siRNA (esiRNA)^50^ was also analyzed. The AS events differentially occurring in each condition (MED13L-, RERE-, or EWSR1::FLI1-knockdown *vs*. control) were quantified according to the percentage of spliced-in (PSI). Classification of splicing categories showed that both MED13L and RERE knockdown specifically increased events belonging to the category of exons and had limited effects on other splicing categories (**Figure 5C**). Prediction of the effects on protein diversity due to inclusion and skipping events favored by MED13L and RERE knockdowns involved alternative protein isoforms and ORF disruption (**Figure S5C**). Importantly, the splicing profiles of MED13L-and RERE-depleted cells were different from those of EWSR1::FLI1-depleted cells, whose downregulation equally affected all the splicing categories (**Figure 5C**), suggesting that the oncogene regulates specific AS subtypes through MED13L and RERE. Validation of some of these changes was performed by RT-PCR in A673 depleted from MED13L and RERE. For instance, genes such as *APAF1*, *BRD8*, *EZH2* and *U2SURP* transcripts underwent exon inclusion, while in genes like *BIN1*, *CLASP1*, *MICAL3*, and *SOX2* exons were excluded (**Figure 5D**). Furthermore, changes in the splicing of these genes could also be observed in other ES cell lines upon MED13L and RERE knockdown (**Figures S5A, S5B**).

GO analysis of the genes with an altered AS in MED13L-and RERE-depleted A673 cells showed that they were involved in chromosome and chromatin organization, histone modifications, organelle regulation and cell cycle processes, and in RERE-depleted cells also macroautophagy processes (**Figure 5E**). The same analysis for genes affected by AS in EWSR1::FLI1-depleted cells showed a functional enrichment in DNA repair, cytoskeleton organization, histone modification, neuronal development and protein modification categories. Therefore, the functional profile of the genes affected by splicing regulation by MED13L and RERE differs from that of the oncogene, since the latter affects genes with a wide range of functions.

To investigate the mechanisms by which MED13L and RERE alter the splicing of the affected genes, we addressed the possibility that MED13L and RERE might directly bind to them and co-transcriptionally regulate their splicing. We crossed the coordinates of the AS events in MED13L-and RERE-depleted cells with the binding sites of MED13L and RERE in the parental A673 cells, previously determined by ChIP-seq, and found that, of approximately 1,500 AS regulated events, MED13L and RERE were bound at the exact same splicing event only in few genes like *ADGRL1* and *RAB11FIP3*. This low percentage of genes with AS and binding of MED13L and RERE indicated that the AS effects observed in MED13L-and RERE-depleted cells were due to the observed reduced expression of various splicing factors in those cells. Consistent with this possibility, extraction of the most prominent genes associated with the biological process of “RNA splicing regulation”, identified by GSEA in our RNA-seq data, revealed widespread downregulation of these genes following MED13L and RERE depletion (**Figure 5F**). These results were validated by RT-qPCR for selected genes in different ES cell lines, and by Western blot in A673 cells (**Figure S5D** and **S5E**).

In addition to several proteins, post-transcriptional regulation of gene expression also requires the action of non-protein-coding factors, and MED13L, RERE and EWSR1::FLI1 bind to the genes encoding certain transcripts involved in RNA metabolism and ribosomal processes, such as the splicing factor *U2* and the ribosomal subunits *45S RNA* and *5.8S RNA* (see above). As the expression of these genes could not be detected by RNA-sequencing of poly-A transcripts, we analyzed their expression levels in Ewing sarcoma cells depleted from FLI1, MED13L or RERE. This approach showed that downregulation of any of these proteins in different ES cell lines resulted in reduced levels of 45S RNA, 5.8S RNA and U2 (**Figures 5G, S5F,** and **S5G**), indicating direct co-regulation of the expression of key genes for AS and translation by Mediator, RERE and EWSR1::FLI1.

### The RBM39 degrader indisulam shows potent antitumor activity in Ewing sarcoma

Direct regulation by the oncogene of MED13L and RERE expression levels and chromatin recruitment suggested that strict control of splicing mediated by these two proteins is essential for ES tumorigenesis. Heatmap representation of the altered expression of genes involved in RNA splicing in A673 cells depleted of MED13L and RERE shows that RBM39 is the most downregulated gene (**Figures 5F**). Furthermore, depletion of these two proteins in other ES cell lines also resulted in reduced RBM39 expression levels (**Figures S5D, S5E**). Since the *RBM39* locus lacks MED13L and RERE binding in A673 cells (**Figure S6A**), these findings suggest transcriptional regulation by distal super-enhancers, although an indirect effect of these proteins on RBM39 transcriptional regulation cannot be excluded.

The splicing factor RBM39 is an RNA-binding protein with key roles in transcriptional regulation and AS and is known to interact with the U2-associated splicing factor SF3B1^51^. Importantly, RBM39 has been identified as the molecular target of the anti-cancer aryl sulfonamide indisulam, that functions as a molecular glue facilitating RBM39 recruitment by DCAF15, a substrate receptor of the CUL4-DDB1-DDA1 E3 ubiquitin ligase complex. As a result of this targeting, RBM39 gets polyubiquitinated followed by proteasome-mediated degradation^51, 52^. Indisulam was originally discovered as a cell cycle inhibitor^53, 54^, as it reduces the expression of proteins involved in the cell cycle by RBM39-mediated AS. Among these, the cyclin-dependent kinase CDK4, which is involved in mitosis, undergoes exon skipping in the neuroblastoma cell lines IMR-332 and KELLY treated with indisulam^55^. Analysis of the AS that leads to reduced CDK4 protein levels in neuroblastoma revealed that this same mis-splicing also occurs in A673 cells treated with indisulam, suggesting that this drug might contribute similarly to DCAF15-dependent cell cycle dysregulation in ES (**Figure S6B**).

In MYC-driven neuroblastoma models, the antitumoral activity of indisulam correlates with the expression levels of RBM39 and DCAF15, and RBM39 has been suggested as a direct target of MYC^55, 56^. Expression data of pediatric tumors from our institution revealed that RBM39 and DCAF15 expression was significantly higher in ES tumors than in other tumors, including neuroblastoma (**Figure 6A**). Visualization of the oncogene binding sites in A673 cells with the UCSC Genome Browser shows that EWSR1::FLI1^11^ binds to both the *RBM39* and *DCAF15* promoter regions (**Figure 6B**). Accordingly, the expression of these two proteins was reduced following inducible EWR1::FLI1 knockdown (**Figures 6C** and **S6C**), indicating that RBM39 and DCAF15 are direct transcriptional targets of EWSR1::FLI1. Immunohistochemical analysis of representative histological sections of ES primary tumors (n=13) showed a high and homogenous expression of RBM39 and DCAF15 in most ES tumors regardless of their clinical stage or chromosomal translocation (**Figure 6D**). Overall, 70% of the tumors presented positive staining. Of note, the tumor sample with negative RBM39 staining also displayed low DCAF15 expression (sample ES3, **Figure 6D**). Expression of RBM39 and DCAF15 was further evaluated in a panel of ES cell lines (A673, A4573, CHLA-9, CHLA-25, COG-E-352, TC205, TC71 and SK-ES1) and compared to rhabdomyosarcoma (RH4) and neuroblastoma cell lines (LAN1 and SK-N-SH) (**Figure 6E**). Remarkably, DCAF15 expression levels were similar-and in many cases higher-in ES cells than in the *MYCN*-amplified LAN1 neuroblastoma cell line, apart from the ES cell line SK-ES1, which had low DCAF15 protein levels. Regarding RBM39, its expression is heterogeneous across all cell lines regardless of tumor type, with several ES cell lines showing the highest levels of RBM39 expression.

**Figure 6.**
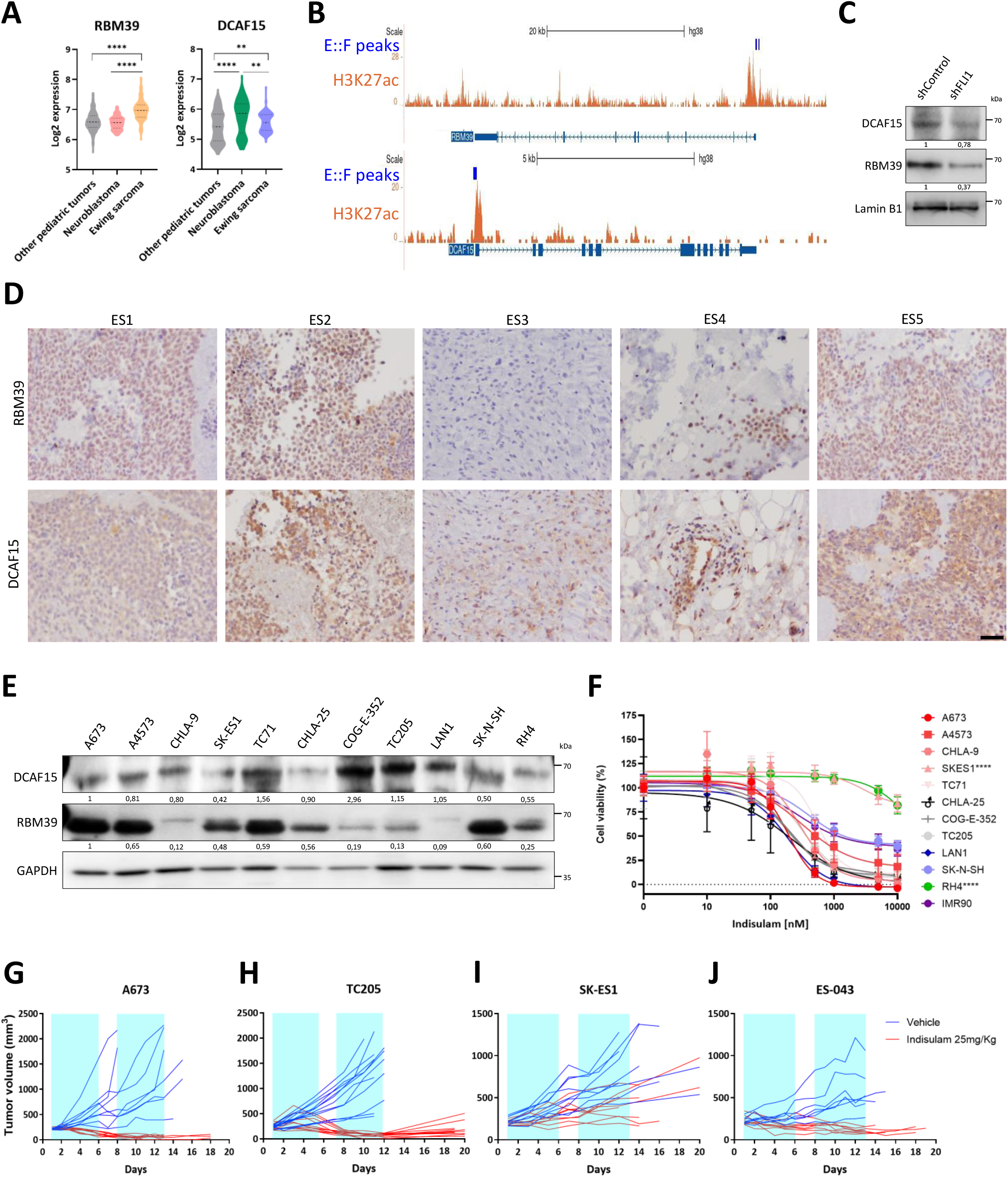
Ewing sarcoma is highly sensitive to the RBM39 degrader indisulam. **A)** mRNA expression of RBM39 and DCAF15 in pediatric tumors (n=519) versus neuroblastoma (n=122) and Ewing sarcoma tumors (n=142), extracted from Hospital Sant Joan de Déu Barcelona DAIOMICS database. Ordinary one-way ANOVA with Tukey’s multiple comparisons test was performed to evaluate the differences. **P <0.01, ****P <0.0001. **B)** UCSC Genome Browser tracks showing enrichment in H3K27ac and EWSR1::FLI111 at RBM39 (chromosome 20) and DCAF15 (chromosome 19) genomic regions in the A673 cell line. E::F, EWSR1::FLI1. **C)** DCAF15 and RBM39 protein levels in A673 cells after induction of EWSR1::FLI1 knockdown with 1 μg/mL doxycycline, determined by Western blot. Lamin B1, loading control. Band intensities were quantified relative to Lamin B1 using ImageJ. **D)** RBM39 and DCAF15 expression in representative tumor samples of Ewing sarcoma obtained from Hospital Sant Joan de Déu Biobank (n=13), detected by IHC. ES1 originated from a locally relapsed tumor, ES2 and ES3 from metastatic tumors and ES4 and ES5 from tumors at diagnosis. All of them have EWSR1::FLI1, except for ES5, which has the FUS::ERG translocation Scale bar: 200 μm. **E)** Protein expression of DCAF15 and RBM39 in the cell lines A673, A4573, CHLA-9, SK-ES1, TC71, CHLA-25, COG-E-352 and TC205 (Ewing sarcoma, LAN1 and SK-N-SH (neuroblastoma) and RH4 (rhabdomyosarcoma). GAPDH, loading control. Band intensities were quantified relative to GAPDH using ImageJ. **F)** Cell viability curves for different cell lines after 72 hours treatment with different concentrations of indisulam. IMR90, fibroblast cell line derived from normal lung tissue. Ordinary one-way ANOVA and Dunnett’s multiple comparisons test were used to compare treatment differences between A673 and the other cell lines. ****P value <0.0001. **G-J)** Progression of the tumor volumes of A673 **(G;** control group: n=9; indisulam group: n=9), TC205 (**H**; control group: n=12; indisulam group: n=11), SK-ES1 (**I**; control group: n=9; indisulam group: n=9), and ES-043 (**J**; control group: n=7; indisulam group: n=8) xenografts treated with 25mg/kg indisulam or vehicle, 5 days on-blue-, 2 days off-white-, for two cycles.

Since RBM39 expression is regulated directly by the oncogene and indirectly through MED13L and RERE in ES cells, we investigated the sensitivity of the different ES cell lines to DCAF15-mediated proteosomal degradation of RBM39. *In vitro* viability assays showed that ES cell lines with high DCAF15 levels (i.e. A673, A4573, CHLA-9 and TC71) had greater sensitivity to indisulam compared to RH4 cells or the fibroblast cell line IMR90, regardless of RBM39 levels (**Figures 6F**). Sensitivity to indisulam was not limited to EWSR1::FLI1-bearing cell lines, since the cell line TC205, which harbors the *EWSR1::FEV* fusion, and the cell lines CHLA-25 and COG-E-352, with the *EWSR1::ERG* fusion, also showed low IC_50_ for indisulam, in line with their high DCAF15 expression levels. In contrast, SK-ES1 cells, which display low DCAF15 expression levels were insensitive to indisulam. Among the neuroblastoma cell models, the MYC-amplified cell line LAN1 had similar sensitivity to indisulam as most ES cell lines, whereas the SK-N-SH cell line without MYCN amplification showed reduced sensitivity compared to ES cell lines.

To further demonstrate the sensitivity of ES cells to indisulam, we evaluated its therapeutic efficacy *in vivo* using xenograft models. To this end, A673 cells were subcutaneously inoculated in mice and, once tumors reached an average volume of approximately 200 mm^3^, mice were treated intraperitoneally with indisulam at 25 mg/kg for two cycles of 5 days on - 2 days off, each. Of note, this drug schedule was well tolerated, and no relevant toxicities were observed in the animals receiving treatment under these conditions (**Figure S6D**). Importantly, a marked reduction in tumor volume was rapidly observed within few days after treatment initiation, and complete tumor regressions were achieved in most tumors (**Figure 6G**). The antitumoral effect of indisulam was also observed in xenografts from TC205 cells, harboring the *EWSR1::FEV* chromosomal translocation (**Figure 6H, S6E**). In contrast, SK-ES1 xenografts model did not respond to indisulam treatment, which is consistent with its low levels of DCAF15 (**Figure 6I, S6F**). Furthermore, the therapeutic efficacy of indisulam was also observed in cells from a patient-derived xenograft (**Figure 6J, S6G**). Altogether, these results demonstrate the extraordinary sensitivity of most ES tumors to RBM39 degrader indisulam and support the use of DCAF15 expression as a biomarker of response.

## DISCUSSION

EWSR1::FLI1 rewires the epigenetic landscape of the cell^3, 10, 11, 17^ and the critical role of the epigenetic context for tumorigenesis was clearly shown in our recently published data demonstrating that the sole expression of the oncogene is sufficient to recreate ES when expressed in human embryonic mesenchymal stem cells^5^. Changes in chromatin conformation, orchestrated by the oncogene, contribute to generate *de novo* enhancers and super-enhancers that, as in many other types of tumors, regulate the expression of genes important for cell state maintenance and tumorigenesis ^11, 37, 57^. In this study, we identified MED13L and RERE as EWSR1::FLI1 super-enhancer-bound targets in the A673 cell line^11^, and confirmed that these two proteins are differentially and highly expressed in ES compared to other developmental tumors.

Consistent with its identification as an EWSR1::FLI1-bound target, RERE mRNA and protein levels are directly dependent on EWSR1::FLI1 expression. In contrast, while MED13L protein levels decrease following EWSR1::FLI1 depletion, MED13L mRNA levels remained constant. This discrepancy between MED13L mRNA and protein levels suggests additional post-translational regulatory mechanisms of MED13L driven by EWSR1::FLI1. For example, the oncogene may regulate the expression of microRNAs or long non-coding RNAs that might interfere with MED13L translation, similar to how miR-30 interacts with the 3’UTR region of CD99 to regulate protein expression without affecting mRNA levels^58^.

We demonstrate that, in ES cells, MED13L (and other Mediator complex subunits), RERE and STAG2 bind to overlapping regions across the genome, mainly in distal intergenic regions. These results suggest that these proteins cooperate to regulate chromatin functions. The interaction between RERE and the Mediator complex or STAG2 had not been reported previously in ES or any other cellular context. Although the Mediator complex is conserved in eukaryotes and mostly functions as a transcription coordinator, subunit composition and functions can differ depending on the cell context ^9, 19, 20, 28^. In line with this, our ChIP-seq results indicate a different and specific role for MED13L and RERE in ES, as supported by their genomic distribution and discrete overlap with EWSR1::FLI1-bound regions, which are not the canonical GGAA regulatory microsatellites.

Genome-wide analysis of the DNA binding motifs for MED13L, RERE and STAG2 showed enrichment in CA microsatellites, particularly in intragenic regions. Hui *et al.* demonstrated that CA microsatellites enhance intron splicing through length-dependent binding of the splicing factor hnRNP-L to CA repeats rich genes such as *eNOS*^59^. Considering the role of CA microsatellites in splicing regulation, the enrichment of MED13L and RERE binding in these repeat elements could explain, in part, the altered pattern of splicing events observed upon MED13L and RERE depletion. To fully address this issue, further experiments to document changes in the AS pattern using different lengths of CA repeats in spliceosome encoding genes and in genes with altered AS by MED13L and RERE should be performed.

Interestingly, MED13L, RERE and STAG2 binding motifs in distal intergenic regions correspond to the TEAD binding motif GGAAT^44^. Noorizadeh *et al.* described that IGF1, an autocrine growth factor for ES, induces the expression of YAP1, which cooperates with TEAD transcription factors to activate genes located at TEAD binding sites, and promoting EWSR1::FLI1 transformation during the early steps of tumorigenesis^60^. Indeed, the changes in MED13L recruitment to chromatin induced by EWSR1::FLI1 in heMSCs suggest that MED13L (and, likely, RERE and STAG2) plays a key role in the multilayered reprogramming of protein expression induced by the oncogene during the early stages of transformation.

Our ChIP-seq experiments revealed that MED13L, RERE and STAG2 overlap with EWSR1::FLI1 in restricted regions of the genome. Immunoprecipitation and immunofluorescence analyses showed that EWSR1::FLI1 interacts with STAG2 and MED13L and co-localizes in the same regions. In addition, RERE and STAG2 colocalized in A673 cell nuclei. Taken together, our results suggest that MED13L, RERE and STAG2 cooperate with EWSR1::FLI1 to establish and maintain chromatin structures that enable the interaction of regulatory elements and their target promoters in ES. This possibility is further supported by studies of STAG2 knockout in ES cell lines, which show how enhancer-promoter interactions were disrupted, inducing a different EWSR1::FLI1 transcriptome that promoted metastatic abilities^47, 48^. To better understand the role of MED13L and RERE in the chromatin architecture of ES and their relationship with STAG2, three-dimensional chromatin conformation experiments are ongoing.

The chromatin regions concomitantly bound by EWSR1::FLI1 and MED13L and RERE are enriched in CpG islands, in many cases display peaks in tandem, and encode non-protein coding RNAs (ncRNAs). It should be noted that these tandem regions did not correspond to the “black” regions annotated in the ENCODE project^61^, except for few regions of ribosomal subunits, indicating that genomic regions enriched with overlapping EWSR1::FLI1, MED13L and RERE were accurately determined.

CpG islands are usually devoid of DNA methylation and often correspond to transcriptionally active regions in promoters^62^. Apart from this, CpG islands are also found in distant regions from annotated transcription start sites. Therefore, CpG islands can be found in ncRNA transcribed regions^62^. In agreement with this, the binding of EWSR1::FLI1, MED13L and RERE to CpG-rich sites could be potentially activating the expression of important ncRNAs for tumorigenesis. This possibility is supported by the enrichment of the transcriptionally activated mark H3K27ac in these regions. Although our experimental approach (sequencing of transcripts with poly-A tails) precludes the detection of ncRNAs, our results demonstrate that EWSR1::FLI1 orchestrates the transcriptional regulation of non-protein-coding genes through Mediator and RERE. Some of these non-protein-coding genes are tRNAs and the ribosomal subunits RNA5S and RNA5-8S. Dysregulation of ribosome biogenesis in cancer not only ensures the protein levels needed to meet high metabolic demands but also facilitates cancer progression by maintaining stem-cell features^63–67^. Our ChIP-seq data and the observed downregulation of genes involved in the ribonucleoprotein complex and ribosome biogenesis following depletion of MED13L and RERE demonstrate that both proteins contribute to the translational rewiring required for ES tumorigenesis.

Despite numerous attempts to generate ES cell lines with complete knockouts of MED13L and RERE with the CRISPR/Cas9 technology, our efforts were unsuccessful, likely due to the impaired expression of proteins involved in RNA metabolism, particular those required for the AS process, such as RBM39, which are essential for the survival of the ES cells. Altered splicing has been described in many types of cancer. Indeed, altered transcription of different splicing factors in cancer cells has been previously reported^68^, and it is well known that EWSR1::FLI1 plays a role in AS by interacting with the splicing machinery and regulating the splicing of important genes^3, 22,27^. In addition, the Mediator complex is also involved in the regulation of AS in other cellular contexts^19, 20^. Our data here shows that EWSR1::FLI1, Mediator and RERE bind to tandemly arrayed chromatin regions of the leading splicing factor U2. We have also shown that the expression of spliceosome components decreases following MED13L and RERE depletion, leading to changes in the splicing pattern in ES cells. Importantly, our bioinformatics analysis revealed that the AS profile induced by EWSR1::FLI1 is different from that regulated by MED13L and RERE. Overall, our data suggests that the oncogene elicits a broad regulation of AS and, by regulating MED13L and RERE expression, specifically controls one type of splicing in a particular subset of genes. Therefore, and in accordance with other groups that have previously suggested the presence of transcription factors that could be managing the utilization of specific splicing sites^69, 70^, we propose MED13L and RERE as mediators of some of the AS events observed in ES.

Given the altered splicing patterns observed in many cancers, the targeting of the splicing machinery has emerged as a potential new therapeutic strategy^71^. Our results identified the splicing factor RBM39, which is significantly downregulated after MED13L and RERE knockdown, as a direct target of EWSR1::FLI1. The aryl sulfonamide indisulam mediates proteosome-dependent degradation of RBM39 by binding RBM39 specifically to the substrate receptor of the CRL4 E3 ubiquitin ligase DCAF15. Indisulam has been used *in vitro* and *in vivo* achieving very good results in myeloma, head and neck cancer, acute myeloblastic leukemia and in *MYCN*-amplified neuroblastoma^55, 56, 72–74^. Our findings support and extend the previously reported *in vitro* and *in vivo* efficacy of indisulam using the A673 Ewing sarcoma model to target the Fanconi anemia/BRCA pathway^75^. By evaluating multiple ES models, we demonstrate the functional relevance of splicing regulation as a targetable vulnerability in ES. Notably, ES cell lines are as sensitive to the *in vitro* antiproliferative effects of indisulam as *MYCN*-amplified neuroblastoma cells, which have been previously described as extremely sensitive^55, 56^.

Corroborating the reported sensitivity to indisulam as a dependency on high levels of DCAF15, SK-ES1 cells show reduced DCAF15 expression levels and is the only ES cell line tested that is insensitive to the anti-proliferative effects of indisulam. While promising, the *in vitro* results were confirmed with clear *in vivo* results showing an extraordinary antitumor effect. Treatment of mice bearing A673 xenografts with indisulam resulted in complete tumor regression. Importantly, the antitumor effects were already detectable within few days of starting treatment and persisted after treatment cessation, indicating the extreme sensitivity of ES to RBM39 degraders. As reported in neuroblastoma, and opposed to all other previous preclinical models tested, we observe complete tumor regressions with remission after cessation of therapy. These strong anti-tumor effects are likely related to RBM39 being involved in several critical features of ES cells like aberrant DNA repair, splicing abnormalities and metabolism rewiring^55, 75, 76^. These results demonstrate that, in addition to *MYCN*-amplified neuroblastoma, Ewing sarcoma is particularly sensitive to RBM39 degraders, and the two entities would be the best cancer types where to test single agent activity in clinical trials. Importantly, phase I and II clinical trials evaluating indisulam in humans with acute myeloid leukemia and different solid tumors, including melanoma, gastric cancer, breast cancer, renal cell carcinoma or colorectal cancer, have not reported significant toxicities or severe adverse events^32^. Therefore, we propose indisulam as a promising new therapeutic strategy in ES using expression levels of DCAF15 as biomarkers for selecting ES patients with the best chances to respond.

## METHODS

### Cell culture

Commercial cell lines were grown in standard conditions. Ewing sarcoma cell lines were obtained from the American Type Culture collection (ATCC) (A673, A4573, SK-ES1 and TC71) and from the Alex’s Lemonade Stand Foundation for Childhood Cancer (CHLA-9, CHLA-25, COG-E-352 and TC205). The rhabdomyosarcoma RH4 and neuroblastoma LAN1 cell lines were purchased from ATCC. HEK-293FT cells, the neuroblastoma cell line SK-N-SH and the fibroblast cell line IMR90 were obtained from “Banc de línies cel·lulars tumorals” of Hospital del Mar Research Institute (Barcelona). A673 TR shFLI1 cell line was obtained from Dr. Javier Alonso’s group^41^ and was treated with 1 µg/mL of Doxycycline hyclate (Sigma Aldrich) during a minimum of 72 hours to knockdown EWSR1::FLI1. heMSCs were grown in Eagle’s medium supplemented with 10% FBS and bFGF (1 ng/ml)^5^.

### Transfection and infection of Ewing sarcoma cell lines

To express a transient knockdown, following the manufacturer’s instructions, Ewing sarcoma cells were transfected with DharmaFECT 4 (Horizon) using 1 µM of siRNA oligonucleotides in serum-free OPTIMEM medium (Gibco). GFP siRNA was used as control and three different siRNA sequences were used to target MED13L and RERE (see sequences in **Table 1**). The transfection process was repeated 24 hours later and cells were collected 72 hours after the first transfection. For stable and inducible knockdown, shRNAs for MED13L and RERE were cloned in-house in the Tet-pLKO-puro construct (SBI System Biosciences) (see sequences in **Table 2**). For ectopic expression of EWSR1::FLI1, FLAG-tagged EWSR1::FLI1 was cloned into the SPARQ vector carrying the GFP reporter gene (SBI System Biosciences). HEK-293FT supernatants of lentiviral viruses expressing shRNAs or the oncogene were collected and used to infect the cells. To select the infected cells with Tet-pLKO-puro vectors, 2 μg/mL of puromycin (Gibco) was added to cell culture 48 hours after infection. To induce the knockdown, cells were treated with 100 ng/mL of Doxycycline hyclate (Sigma Aldrich) for shMED13L and shRERE expression.

### RNA extraction and real time quantitative PCR (RT-qPCR)

RNA was purified using GenElute Mammalian Total RNA Miniprep Kit (Sigma-Aldrich) or Cytiva Illustra RNAspin Mini Isolation Kit (Cytiva) following the manufacturer’s instructions. Purified RNA was retrotranscribed using from 500 to 2000 ng of RNA. Reverse transcription (RT) was performed with hexamers (Roche, 600 µM) or Oligo (dT)s (Roche, 50 µM), Buffer 5X (Roche), deoxynucleotide triphosphates (dNTPs) (Roche, 10 mM), RNase inhibitor (Roche, 40 units/µL) and transcriptase reverse (Roche, 500 units). In addition, GoScript™ Reverse Transcriptase system (Promega) was used in some experiments. cDNA obtained after RT was analyzed by real time quantitative PCR (RT-qPCR) using SYBR Select Master Mix (Applied Biosystems) and specific forward and reverse primers pairs (see **Table 3**) in 7900HT Fast Real-Time PCR (Thermo Fisher Scientific) or in QuantStudio 5 (Thermo Fisher Scientific) in 384-well plates. Analysis of the small nuclear RNA (snRNA) RNU2-1 expression was performed with TaqMan Gene Expression Assay 20X for RNU2-1 (Hs02786874_gH, Applied Biosystems) and TaqMan Universal PCR Master Mix (Applied Biosystems). Final data was analyzed with the fold change method 2^-ΔΔCt^ using Actin B or GAPDH as housekeeping genes.

### Western Blot (WB)

Protein extraction was performed using RIPA-M lysis buffer and Urea-T buffer as previously described^77^. In subcellular fractionation experiments, cells were pelleted and resuspended in Buffer A (0.25 M Sucrose (Sigma-Aldrich), 10 mM HEPES pH 7.5, 3 mM CaCl2 (Sigma-Aldrich), 10 mM NaCl, 1 mM Phenylmethanesulfonyl fluoride (PMSF) (Sigma-Aldrich), 1 mM Dithiothreitol (DTT) (Cytiva), 0.25% NP-40 and 1X Roche’s protease inhibitors). Samples were incubated for 30 minutes at 4^0^C and centrifuged 10 minutes at 3000 rpm at 4^0^C. Supernatants corresponded to the cytoplasmic fraction and were collected. Pellets were rinsed twice with Buffer A and supernatants were collected as cytoplasmic fraction. Nuclear fraction was extracted using RIPA-M and Urea-T buffer, as previously described^78^. 20 to 50 µg of protein extracts were mixed with loading Laemmli buffer to perform WB following standard protocols. Secondary antibodies coupled to horseradish peroxidase (HRP) (Dako) were used and light signals produced by Immobilon ECL Ultra Western HRP Substrate (Merck Millipore) reacting to HRP were detected in a dark room using SRX-101A medical film processor (Konica-Minolta) or using Imager Vilber Fusion FX7 (Vilber). Relative protein expression was quantified using ImageJ software. Primary antibodies used were: Actin B, FLI1, Lamin B1, RBM39, U2AF2 (Santa Cruz Biotechnology); DCAF15 (Merck), FLAG, GAPDH (Sigma-Aldrich); MED1, MED12, MED13L (Bethyl); and RERE (Invitrogen).

### Immunohistochemistry (IHC)

Sections from paraffin blocks of tumor tissues from Hospital Sant Joan de Déu Biobank and experimental tumors were analyzed by immunohistochemistry of DCAF15 (Merck), MED1, MED12, MED13L (Bethyl); RERE (Sigma-Aldrich) and RBM39 (Santa Cruz Biotechnology), and counterstained with hematoxylin following standard methods^79^.

### Immunoprecipitation (IP)

A673 cells were lysed with 0.5% Triton X-100, 1 mM EDTA, 1X Roche’s protease inhibitors and 1X Roche’s phosphatase inhibitors. Supernatants were collected and pre-cleared using 0.5% Triton X-100, 1mM EDTA, 1% BSA, 1 µg rabbit IgG, 25 µL of Dynabeads A (Invitrogen) and 25µL of Dynabeads B (Invitrogen), 1X Roche’s protease inhibitors and 1X Roche’s phosphatase inhibitors. Then, supernatants were incubated with 25 µL/25 µL of Dynabeads A/ B, 3 µL of the specific antibody (FLAG, RERE (Sigma-Aldrich); FLI1 (Santa Cruz Biotechnology), MED1, MED12 or MED13L (Bethyl)) and lysis buffer at 4°C overnight. IPs with IgG (Abcam) were used as negative controls of protein binding. Beads were resuspended in 1X Laemmli Buffer and boiled to denature proteins and unbind the beads. Beads were discarded and samples were loaded in an 8% SDS-PAGE gel to analyze protein-protein interactions by WB.

### Immunoflourescence (IF)

Cells seeded on glass slides were fixed with 3.7% formaldehyde solution in PBS and permeabilized with 0.2% Triton X-100 in PBS. Non-specific binding was blocked using 1% BSA and 0.03% Tween 20 in 1X PBS and, after 30 minutes, slides were incubated with primary antibodies (FLAG, RERE (Sigma-Aldrich); FLI1, STAG2 (Santa Cruz Biotechnology); and MED13L (Bethyl)). After several washes, samples were incubated with secondary antibodies (Alexa Fluor 546 or Alexa Fluor 647, Invitrogen) in blocking buffer. Cell nuclei were stained with 4′,6-diamidino-2-phenylindole (DAPI) (Invitrogen) before mounting the slides with Fluoromount G (Thermo FIsher). Images corresponding to the different channels were captured with Confocal Multispectral Leica TCS SP8 microscope at the Microscopy Confocal Unit, Hospital Sant Joan de Déu Barcelona. Super-resolution images were obtained by image deconvolution using HyVolution tool from Leica Microsystems and were processed and analyzed with LAS X (Leica) and Fiji/ImageJ softwares. Colocalization experiments were quantified by Overlap Coefficient and Förster resonance energy transfer (FRET) 135 using LAS X software (Leica).

### RNA-sequencing (RNA-seq) and functional analysis

Biological triplicates including three different siRNA sequences targeting MED13L and three different sequences for RERE (see **Table 1**) were used to perform an RNA-seq using poly-A capture method. Samples were sequenced in a HiSeq 2500 platform (Illumina) with a paired-end coverage and ∼30 million reads/sample using a 2 x 75 flowcell sequencer at the Genomics Unit of the Center for Genomic Regulation (CRG, Barcelona). Raw sequencing reads in the fastq files were mapped with STAR version 2.7.1a^80^. Gencode release 36 based on the GRCh38.p13 reference genome was used to annotate transcripts. Genes having less than 10 counts in at least 2 samples were excluded from the analysis. Pre-Ranked Gene Set Enrichment Analysis (GSEA)^81^ implemented in clusterProfiler package (version 4.0.0)^82^ was used in order to retrieve enriched functional pathways. The ranked list of genes was generated using the-log(p.val)*signFC for each gene from the statistics obtained in the DE analysis with limma^83^. Functional annotation was obtained based on the enrichment of gene sets belonging to gene set collections in Molecular Signatures Database (MSigDB). The collections used in this project are c5.bp: Gene sets derived from the Biological Process Gene Ontology (GO) (version 7.2). Data analysis performed with R (version 4.1.0) included limma, genefilter, gplots, Vennerable, and hugene20sttranscriptcluster.db packages.

### ChIP-sequencing, ChIP-qPCR and bioinformatic analysis

Chromatin immunoprecipitation (ChIP) assays using 10 µg of H3K27ac (Abcam), MED1, MED12, MED13L (Bethyl); RERE (Invitrogen) and STAG2 (Santa Cruz Biotechnology) antibodies were performed as described by García-Hernández, *et al*.^84^ or following the manufacturer’s instructions of the ChIP-IT High Sensitivity kit (Active Motif). Sequencing was carried out in a HiSeq 2500 platform (Illumina) with 30 – 40 million reads/sample using a 1 x 50 sequencer. All samples were included in technical triplicates for A673 cell line and duplicates for heMSCs cells. Annotation of the peaks was done using the ChIPseeker package^85^. GENCODE version 39 was used to annotate the peaks using version 3.15.0 from TxDb.Hsapiens.UCSC.hg38.knownGene package. The consensus peak set was obtained by overlapping the three replicates and retaining peaks found in at least two replicates using the function findOverlapsOfPeaks of the ChIPpeakAnno package (version 3.30.1) using different windows in the *maxgap* argument. Consensus Peak Annotation was performed by annotatePeak. Promoter region was defined from 5kb upstream to +100bp downstream of the transcription start site (TSS). The position and strand information of nearest genes were reported, as well as the distance from the peak to the transcription start site of its nearest gene and the genomic region. Since some annotation overlapped, ChIPseeker adopted the following priority in genomic annotation: Promoter > 5’ UTR > 3’ UTR > Exon > Intron > Downstream > Intergenic. Downstream was defined as the downstream of gene end. EWSR1::FLI1 consensus peaks and annotations were analyzed from publicly available raw data of EWSR1::FLI1 ChIP-seq in A673 cell line^43^ at NCBI Sequence Read Archive (SRA; http://www.ncbi.nlm.nih.gov/sra) under accession number SRA096176. DNA binding motif of each consensus peak set and of the overlapping peaks between different consensus peak sets was analyzed using MEME-ChIP web service^86^. The University of California Santa Cruz (UCSC) Genome Browser (https://genome.ucsc.edu/) was used to visualize the genomic data obtained^87^. For ChIP-qPCR, eluted chromatin and input samples were used for RT-qPCR as previously described (see **Table 3** for primer sequences). Data was analyzed calculating the percent of input for each ChIP: %Input = 2^(-ΔCt^ ^[normalized^ ^ChIP])^ x 100 where normalized ChIP is Ct[ChIP] - (Ct[Input] - Log2(Input Dilution Factor)).

### Splicing analysis

RNA-seq data was used to perform a splicing analysis after MED13L and RERE depletion by siRNA. Analysis was carried out by using the pipeline VAST-TOOLS^49^ to calculate the metric percent spliced-in (PSI) and compare the control with the studied condition. A public RNA-seq database of EWSR1::FLI1 knockdown by endoribonuclease-prepared siRNA (esiRNA)^50^ was used. Splicing events were classified into exons, intron retention, alternative 3’ splice site (or alt. acceptor) and alternative 5’ splice site (alt. donor) events. A filter of delta PSI ≥ 15 was applied for each splicing category. The number of events higher in control, higher in MED13L, RERE, or EWSR1::FLI1 upon knockdown was obtained. Results were validated performing a RT-PCR with the retrotranscribed cDNA obtained by siMED13L and siRERE samples (see **Table 4** for primer sequences). RT-PCR products were loaded into 1% agarose or 8% polyacrylamide gels stained with SYBR Safe DNA Gel Stain (Invitrogen). Gel visualization and image capturing was performed by UV light exposure in Gel DocTM XR+ Gel Documentation System (Bio-Rad) using Image LabTM software (Bio-Rad). Analysis of band intensities with ImageJ software allowed to calculate PSI as (inclusion/(inclusion+exclusion)*100.

### Cell viability assays

Cells (3000/well) were seeded in 96-well plates and, 24 hours later, treated with indisulam for 72 hours. Cell viability was assessed with the Luminescence Cell Viability Assay Cell Titer-Glo (Promega). Luminescence signal was detected by using the Orion II luminometer (Berthold) and IC50 calculations were performed using Graphpad Prism 8.0.

### In vivo studies

*In vivo* studies were accomplished according to the institutional and European guidelines (EU Directive 2010/63/EU). Animal procedures have been approved by the animal experimental ethics committee (Comité de Ética de Experimentación Animal, University of Barcelona). 1×10^6^ A673, SK-ES1 and TC205 cells or 3 × 3 mm^3^ fresh PDX tumor from HSJD-ES-043 were subcutaneously inoculated into three-to six-week-old athymic nude mice. Mice were weighed and tumors measured with a caliper three times per week. Tumor volume was calculated as follows: (longer measurement x (shorter measurement)^2)/2. Indisulam was diluted in 10% DMSO and 90% of 20% SBE-β-CD. Once tumors reached an average volume of approximately 200 mm^3^, mice were randomized into indisulam treatment (25 mg/kg) or control vehicle group. Treatment was administrated intraperitoneally five times per week with 2 days off each week, during two consecutive weeks. Animals were sacrificed according to the humane endpoint criteria at the end of the treatment or if tumor volume exceeded 1500 mm^3^.

### Statistical analysis

Statistical analyses were conducted using the software Graphpad Prism 8.0. Data is expressed as mean values ± standard deviation (SD). Tests performed in order to compare two groups are specified in each experiment. Statistical significance was determined by a p value of less than 0.05.

