## Supplementary figures for "RERE and the Mediator complex cooperate with EWSR1::FLI1 in the reprogramming of Translation and Alternative Splicing, the latter being a therapeutically targetable vulnerability in Ewing sarcoma"

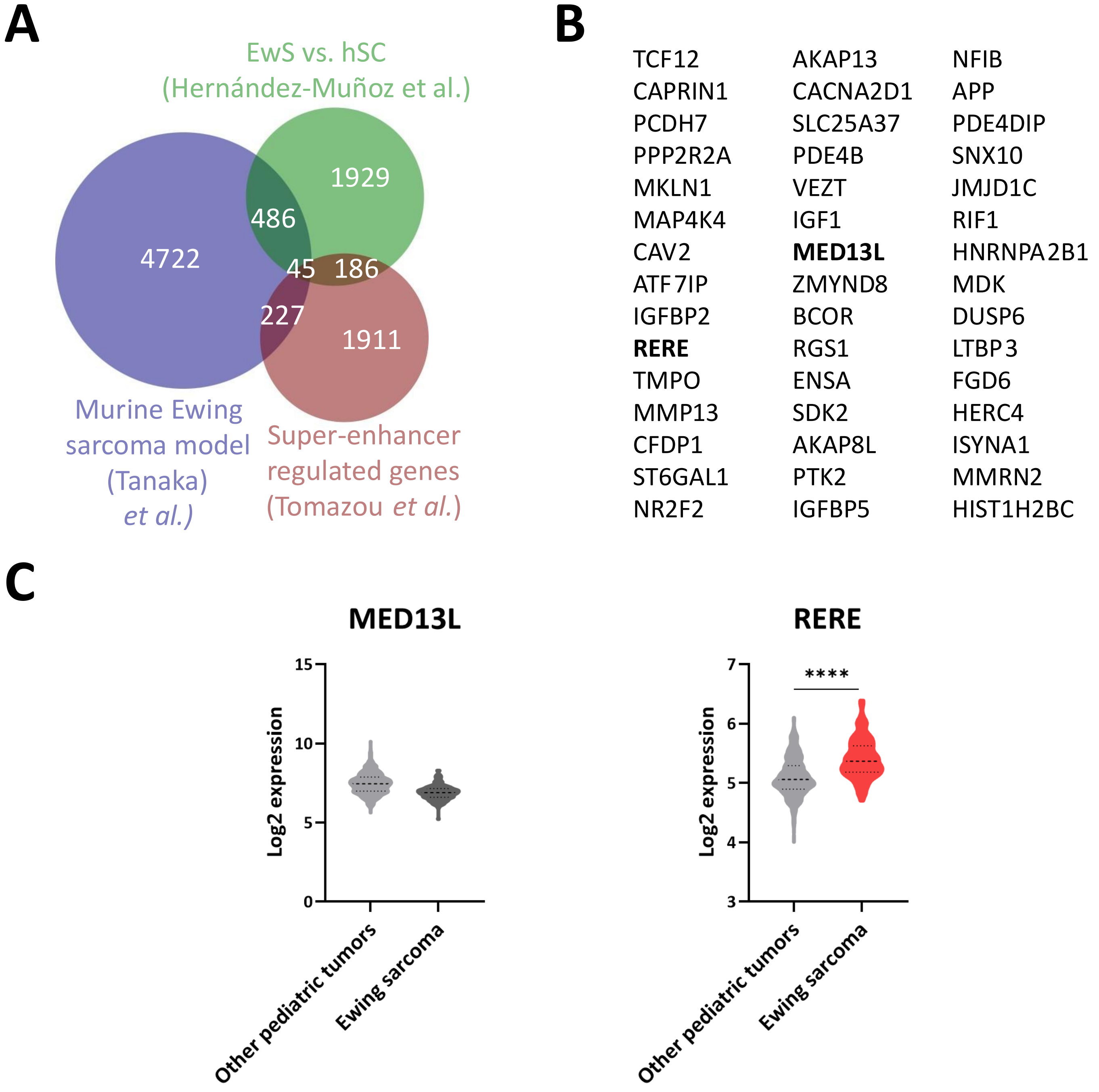


***Supplementary Figure 1. Identification of super-enhancer-regulated genes important for tumorigenesis by in silico data mining. A)*** *Venn diagram depicting the strategy of in silico data mining to identify super-enhancer-bound genes important for Ewing sarcoma tumorigenesis.* ***B)*** *List of the 45 common genes identified through in silico data mining.* ***C)*** *mRNA expression of MED13L and RERE in pediatric tumors (n=641) versus Ewing sarcoma tumors (n=142), extracted from Hospital Sant Joan de Déu Barcelona DAIOMICS database. Unpaired t test by the Mann Whitney test was performed to evaluate the differences. ****P <0.0001.*

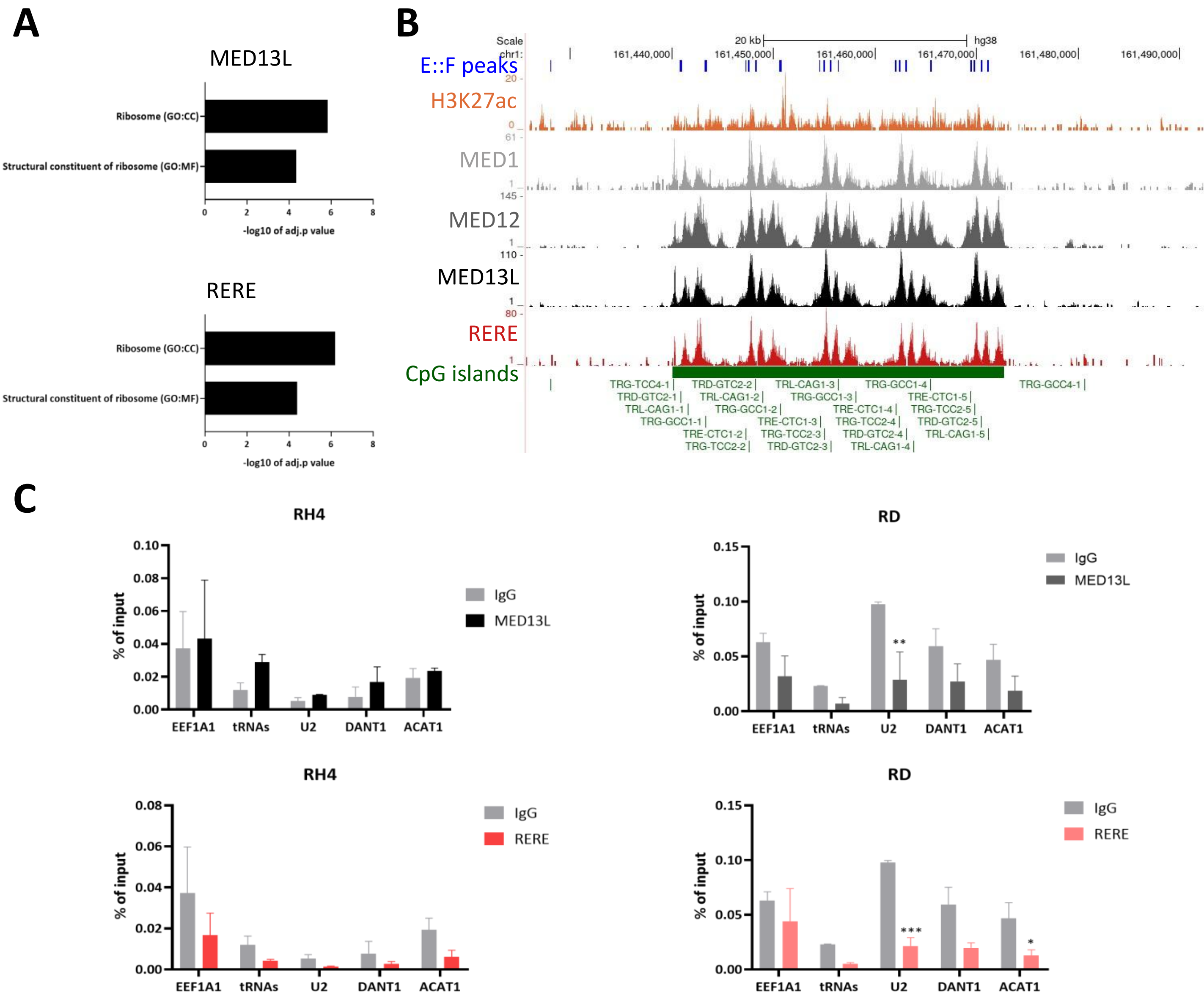


***Supplementary Figure 2. Chromatin binding of MED13L and RERE in Ewing sarcoma and rhabdomyosarcoma cell lines. A)*** *Bar plots displaying statistically significant categories from Gene Ontology (GO) analysis of the genes bound by MED13L, RERE and EWS::FLI1 in A673 cells. Adjusted P value <0.05. Analysis was performed with gProfiler online database[90] (https://biit.cs.ut.ee/gprofiler/gost). CC, cellular component; GO, Gene ontology; MF, molecular function.* ***B)*** *UCSC Genome Browser track showing the enrichment of FLI1[43], H3K27ac, MED1, MED12, MED13L and RERE in tRNA genic regions (chromosome 1). E::F, EWSR1::FLI1.* ***C)*** *MED13L and RERE binding to targets overlapping with EWSR1::FLI1 in the rhabdomyosarcoma cell lines RH4 and RD, determined by ChIP-qPCR. Data represent the enrichment ratio of immunoprecipitated samples relative to input. IgG, negative control for each immunoprecipitation; ACAT1, control region. Two-way ANOVA was used to analyze differences and Sidak's multiple comparisons test was performed to compare enrichment versus IgG. **P value <0.005, ***P value <0.001.*

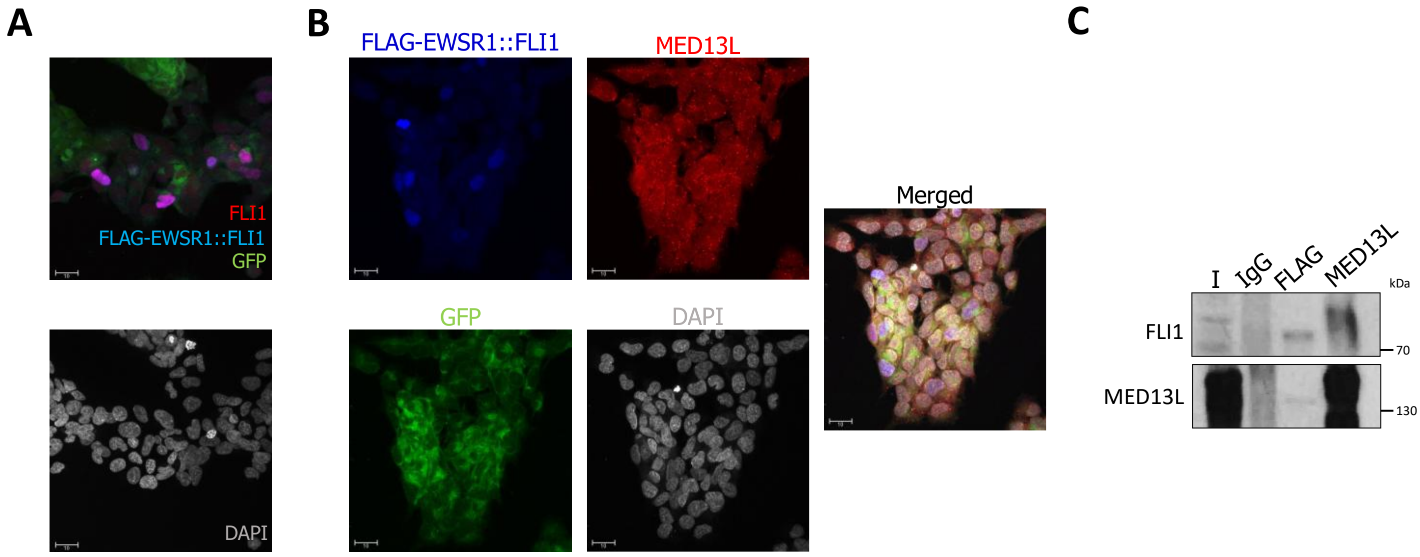


***Supplementary Figure 3. MED13L and STAG2 interact with EWSR1::FLI1 in discrete nuclear domains.*** ***A)*** *A673 cells were infected with lentiviral FLAG-EWSR1::FLI1 supernatants, and double immunofluorescence (IF) to detect FLAG (blue) and FLI1 (red) was performed. Cell nuclei were identified by DAPI staining (grey). GFP (green) indicated cells infected with FLAG-EWSR1::FLI1. Scale bar: 10 µm.* ***B)*** *IF detection of FLAG (blue) and MED13L (red) in A673 cells expressing FLAG-EWSR1::FLI1. Cell nuclei, in grey. Scale bar: 10µm.* ***C)*** *A673 cells were infected with lentiviral FLAG-EWSR1::FLI1 supernatants and cell extracts were generated. Immunoprecipitations using antibodies against FLAG and MED13L were performed, and bound proteins were detected by WB.*

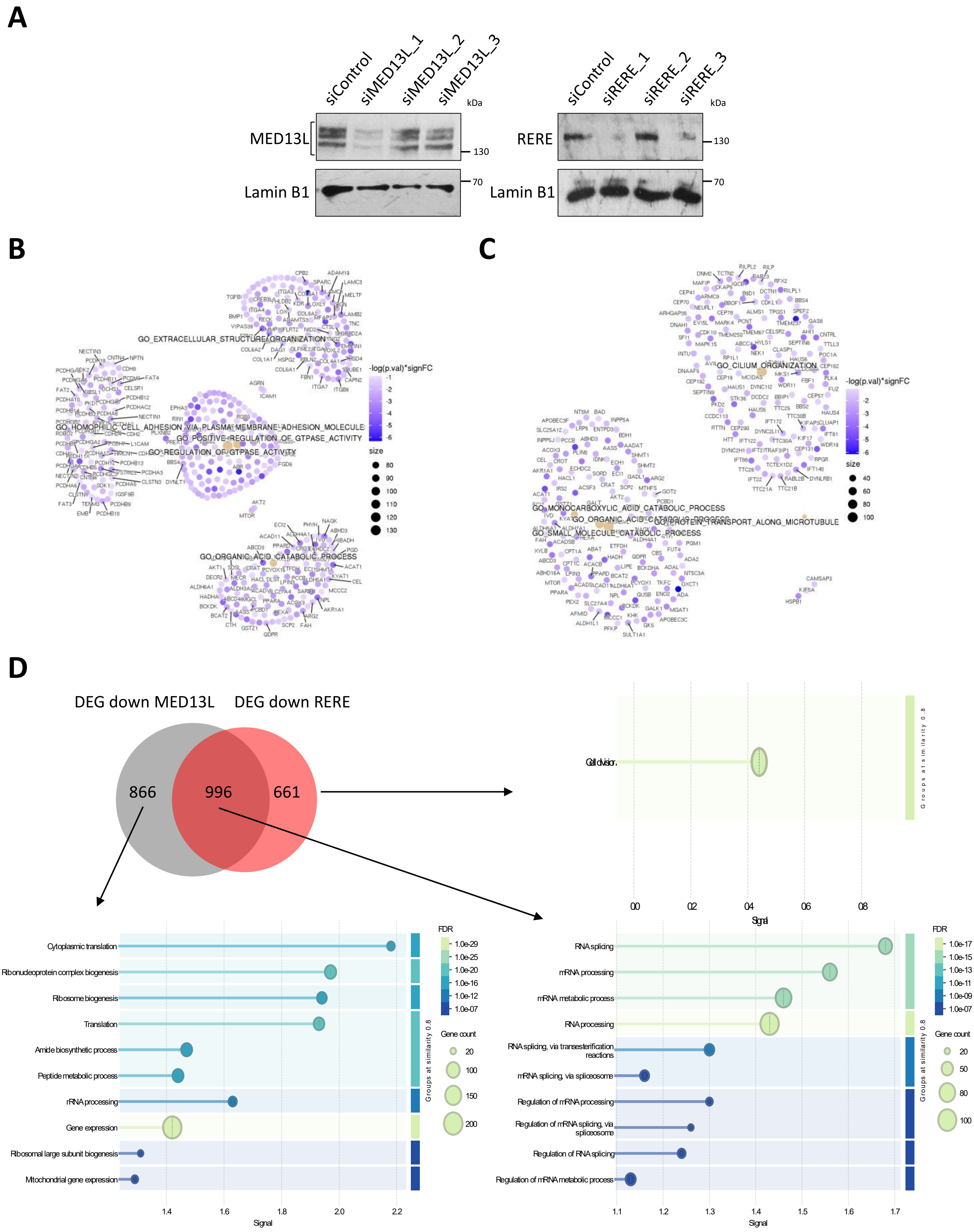


***Supplementary Figure 4. Functional effects of MED13L- and RERE-depleted cells in Ewing sarcoma models. A)*** *Western blot validation of MED13L and RERE depletion using three different siRNA sequences in the A673 cell line. Lamin B1, loading control. Square brackets, protein isoforms detected.* ***B, C)*** *Gene-Concept Networks of the top five significantly upregulated terms in MED13L* ***(B)*** *and RERE* ***(C)*** *depleted cells.* ***D)*** *Overlap of transcripts downregulated in A673 cells depleted from MED13L and RERE, and their corresponding GO performed with STRING[88] ([https://string-db.org/](https://string-db.org/)). FDR, False Discovery Rate. .*

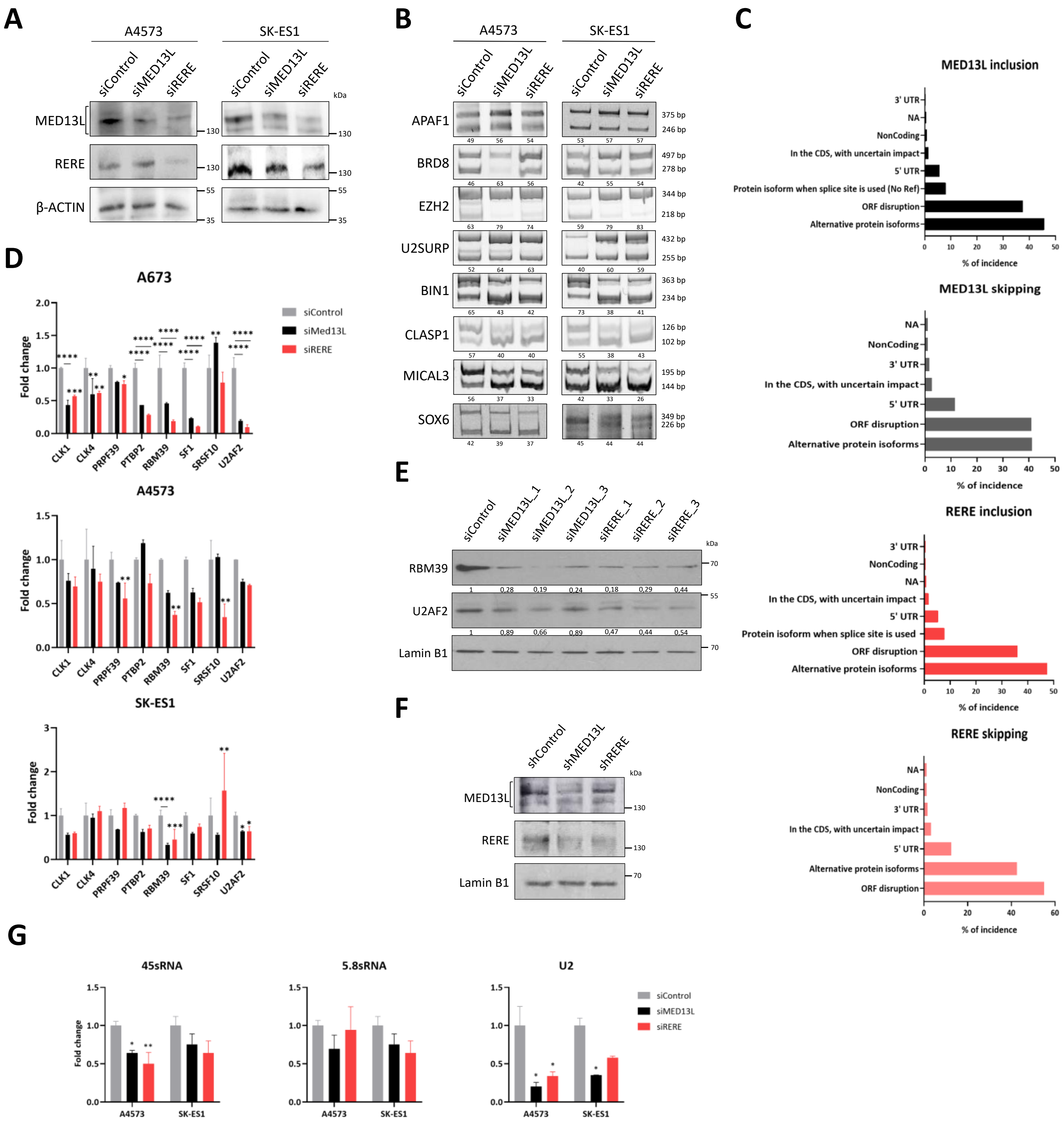

A
A4573
SK-ES1
siControl
siMED13L
siRERE
MED13L
RERE
β-ACTIN
kDa
B
APAF1
BRD8
EZH2
U2SURP
BIN1
CLASP1
MICAL3
SOX6
C
MED13L inclusion
MED13L skipping
RERE inclusion
RERE skipping
3' UTR
NA
NonCoding
In the CDS, with uncertain impact
5' UTR
Protein isoform when splice site is used (No Ref)
ORF disruption
Alternative protein isoforms
% of incidence
D
A673
A4573
SK-ES1
Fold change
siControl
siMed13L
siRERE
CLK1
CLK4
PRPF39
PTBP2
RBM39
SF1
SRSF10
U2AF2
E
siMED13L_1
siMED13L_2
siMED13L_3
siRERE_1
siRERE_2
siRERE_3
RBM39
U2AF2
Lamin B1
F
shControl
shMED13L
shRERE
G
45sRNA
5.8sRNA
U2

***Supplementary Figure 5. Alternative splicing regulation by MED13L and RERE. A)*** *Depletion by siRNA validated by Western blot in A4573 and SK-ES1 cell lines using sequence #1 for MED13L and RERE siRNA knockdown. Actin B, loading controls. Square brackets, protein isoforms detected.* ***B)*** *Alternative splicing changes in selected genes induced by MED13L and RERE depletion by siRNA in A4573 and SK-ES1 cell lines, determined by RT-PCR. Sequences #1 for MED13L and RERE siRNA knockdowns was used to perform these experiments. PSI, calculated with ImageJ for each gene.* ***C)*** *Prediction of the protein impact caused by AS changes in exon inclusion and skipping after MED13L and RERE depletion by siRNA in A673 cells. Protein impact dataset was downloaded from VastDB online database (https://vastdb.crg.eu/). CDS, coding sequence; ORF, open reading frame; UTR, untranslated region.* ***D)*** *mRNA levels of different splicing factors after MED13L and RERE depletion by siRNA in the A673, A4573 and SK-ES1 cell lines. Values were normalized to GAPDH and Actin B. Statistical tests to study significant differences versus the control was performed with two-way ANOVA and Dunnett's multiple comparisons test. *P value <0.05, **P value <0.01, ***P value <0.001 and ****P value <0.0001.* ***E)*** *Changes in protein expression of splicing factors after MED13L and RERE knockdown using three different siRNA sequences in A673 cells, determined by WB. Lamin B1, loading control. Band intensities were quantified relative to Lamin B1 using ImageJ.* ***F)*** *Validation of MED13L and RERE knockdown induced with 100 ng/mL doxycycline in the A673 cell line, determined by WB. Lamin B1, loading control. Square brackets, protein isoforms detected.* ***G)*** *45s and 5.8s ribosomal RNA and U2 snRNA expression in A4573 and SK-ES1 cells depleted from MED13L and RERE. Values were normalized to Actin B. Statistical test to study significant differences was performed with two-way ANOVA and Dunnett's multiple comparisons test. *P value <0.05 and **P value <0.01.*

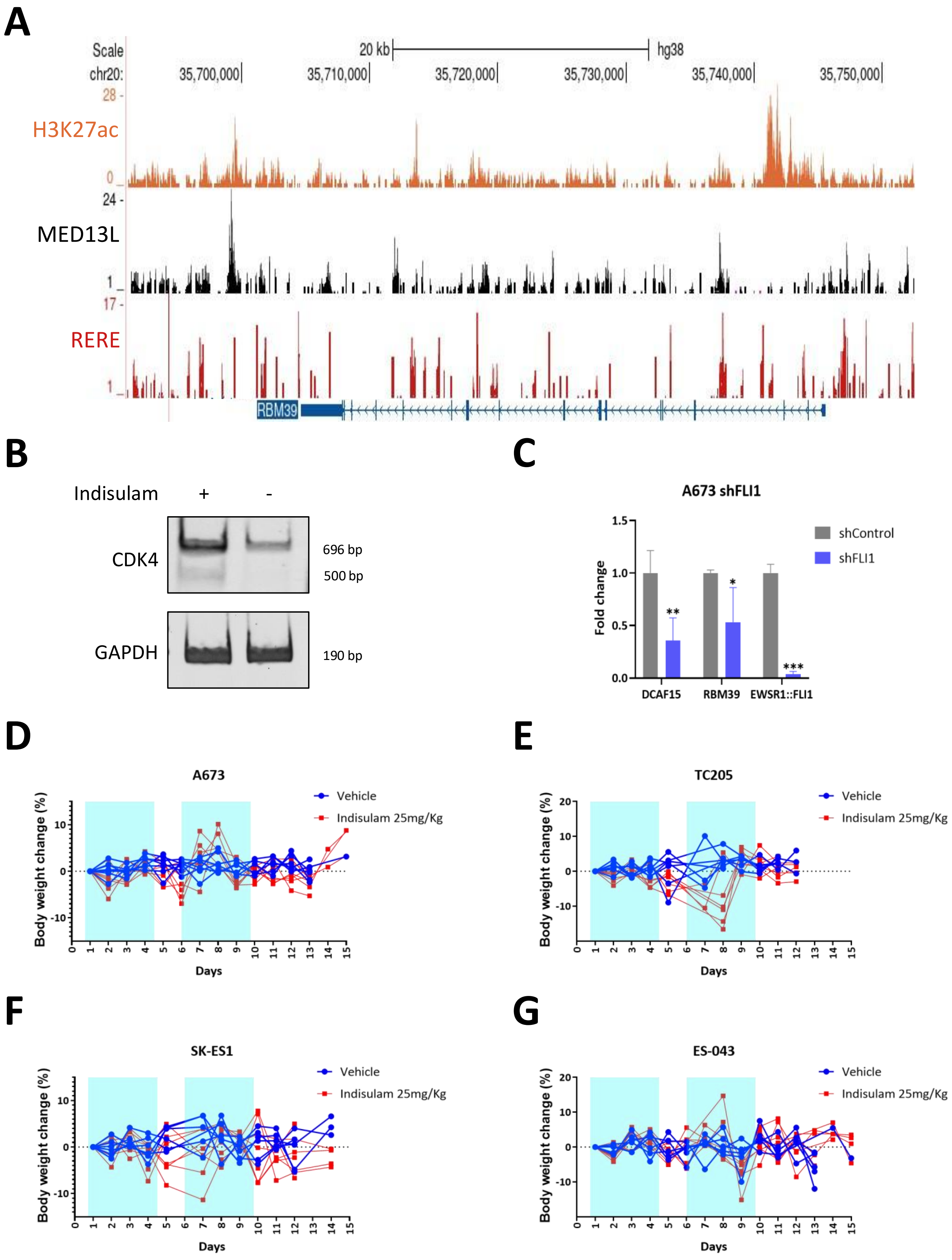


***Supplementary Figure 6. RBM39 and DCAF15 regulation by EWSR1::FLI1 in Ewing sarcoma. A)*** *UCSC Genome Browser tracks for H3K27ac, MED13L and RERE at RBM39 genomic region (chromosome 20) in the A673 cell line.* ***B)*** *CDK4 and GAPDH RT-PCR in A673 cells treated with 190nM indisulam or DMSO for 24 h.* ***C)*** *mRNA levels of DCAF15 and RBM39 in A673 cells after inducible expression of shFLI1 with 1 µg/mL doxycycline, as determined by RT-qPCR. Values were normalized to Actin B. Two-way ANOVA was used to analyze differences and Sidak's multiple comparisons test was performed to compare expression in shFLI1 cells versus control cells. *P value <0.05, **P value <0.01, and ***P value=0.0001.* ***D-G)*** *Progression of the percentage of body weight changes during vehicle and indisulam treatment of mice with A673, TC205, SK-ES1 and ES-043 xenograft models (25mg/kg indisulam, 5 days on -blue-, 2 days off -white-, for two cycles).*
